# ER-Lysosome Cholesterol Exchange Regulates Lysosomal Motility Through mTOR-Dependent LAMTOR1 Phosphorylation

**DOI:** 10.64898/2026.03.31.715514

**Authors:** Pathma Muthukottiappan, Alireza Dehghani, Asisa Muchamedin, Michael Ebner, Mariana E. G. de Araujo, Cristina Coman, Sönke Rudnik, Milash Balachandran, Sofía Fajardo-Callejón, Fatema Akter, Norbert Rösel, Paul Saftig, Markus Damme, Robert Ahrends, Lukas Huber, Volker Haucke, Volkmar Gieselmann, Dominic Winter

## Abstract

The subcellular distribution of lysosomes, the main degradative organelles of mammalian cells, responds to metabolic cues in a highly dynamic way. While lysosomal positioning due to amino acid levels is well-characterized, cholesterol-dependent regulation of lysosomal motility is incompletely understood. We explored impaired lysosomal cholesterol export using a mass spectrometry-based multi-OMICs approach, identifying widespread reallocation of resources and signaling pathway modulation. We identified increased phosphorylation at LAMTOR1 serine 56 in response to cholesterol level perturbations. We demonstrate that this phosphorylation site is sufficient to disrupt Rag GTPases/SLC38A9 binding to the Ragulator complex, inhibiting canonical mTORC1 and facilitating binding of BORC, therefore promoting lysosomal retrograde movement. LAMTOR1 S56 phosphorylation responds exclusively to depletion of lysosomal limiting membrane cholesterol, is facilitated by mTOR, and presents a negative feedback loop for amino acid independent displacement of Ragulator bound Rag GTPases, limiting canonical mTORC1 activity. Mass spectrometry data are available via ProteomeXchange with identifier PXD073489.

**Highlights:**

- Perturbation of lysosomal cholesterol homeostasis results in adaptation of cellular protein and lipid biosynthesis
- LAMTOR1 is phosphorylated at serine 56 via mTORC1
- LAMTOR1 S56 phosphorylation is lysosomal membrane cholesterol dependent
- LAMTOR1 S56 phosphorylation disrupts binding of Rag GTPases to the Ragulator complex
- LAMTOR1 S56 phosphorylation promotes binding of Ragulator to BORC, facilitating lysosomal retrograde transport

## Introduction

Cholesterol is essential for all animal life, providing structural, metabolic, and signaling functions in virtually every cell, and acting as precursor for a variety of biomolecules such as steroid hormones or cholic acids. Disturbed cholesterol homeostasis has been linked to various pathological conditions, such as cardiovascular diseases (e.g. atherosclerosis) or lysosomal storage disorders (e.g. Niemann Pick Type C 1/2 (NPC1/2) deficiency),^1^ and lysosomal dysfunction was shown to strongly alter subcellular cholesterol distribution.^2^

While all cells possess the ability to de novo synthesize cholesterol from acetyl-CoA, they typically acquire it through receptor-mediated endocytosis of low-density lipoprotein (LDL) particles or chylomicron remnants (hepatocytes).^3^ After LDL endocytosis, the receptor dissociates, is recycled, and LDL particles are degraded within lysosomes. This includes liberation of cholesterol from its esterified form by lysosomal acid lipase (LIPA), its sequestering by NPC2, and its export from the lysosomal lumen to the lysosomal limiting membrane either by NPC1 or lysosomal integral membrane protein 2 (SCARB2/LIMP2).^4,5^ Subsequently, cholesterol is transported to the endoplasmic reticulum (ER), the main cholesterol regulatory hub of the cell, through membrane contact sites and several proteins such as vesicle-associated membrane protein-associated proteins A and B (VAPA/B), oxysterol-binding proteins (OSBPs), and OSBP-related protein 1L (ORP1L).^6^ Importantly, cholesterol levels are sensed at the ER by sterol regulatory element-binding proteins 1 and 2 (SREBP1/2). Low ER cholesterol levels result in SREBP transition to the Golgi Apparatus and its proteolytic cleavage, nuclear translocation and an immediate transcriptional response to increase cholesterol de novo synthesis. With approx. 100 ATP equivalents per cholesterol molecule, this process requires a substantial energy input, presenting one of the most resource intensive biosynthetic reactions for a small molecule known for mammalian cells.^7^ Lysosomal cholesterol salvage is, therefore, crucial for cellular energy homeostasis. Despite the important role lysosomes play in this context, the regulatory mechanisms governing their cholesterol handling remain incompletely understood.

Considering cholesterol’s high energetic demand, it is not surprising that its availability influences signaling pathways controlling cellular metabolism, such as the mechanistic target of rapamycin (mTOR),^8,9^ a central metabolic regulator that functions in two distinct complexes, mTORC1 and mTORC2.^10^ Nutrient-dependent signaling promotes the recruitment of mTORC1 to the lysosomal membrane, where it becomes activated and integrates metabolic inputs from amino acids, cholesterol, and others. This promotes anabolic processes, while suppressing catabolic pathways such as autophagy.^11–13^ Crucial components in this process are the heteropentameric Ragulator complex, consisting of late endosomal/lysosomal adaptor and MAPK and MTOR activator (LAMTOR) 1-5, and the Ras-related guanosine triphosphate (GTP)-binding (Rag) GTPases, whose GTP loading state is crucial for the recruitment and activation of mTORC1. For both amino acid and cholesterol levels, direct regulation of mTORC1 through modulation of Rag GTPase activity has been demonstrated,^14^ and it has further been shown that the interaction between Rag GTPases and the Ragulator complex can act as a self-limiting mechanism to restrict mTORC1 activity over time.^15^

Nutrient availability was also shown to strongly influence lysosomal spatial organization. Under conditions of nutrient-sufficiency, lysosomes are predominantly small, highly motile, and localized at the cellular periphery, promoting mTORC1 activity and, hence, signaling. In contrast, nutrient depletion leads to enlarged more static lysosomes that cluster in the perinuclear region. These lysosomes are primarily degradative, harbor inactive mTORC1, and frequently engage in ER contact sites to facilitate metabolite exchange.^8,16^ Importantly, lysosomal mis-localization has been reported for ∼80 pathological conditions.^17,18^ The mechanisms controlling lysosomal positioning are highly complex and only partially understood. At least five protein transport machineries (dynein-dynactin, ADP-ribosylation factor-like protein 8B (ARL8B), Pleckstrin homology domain-containing family M member 2 (SKIP), C-Jun-amino-terminal kinase-interacting protein 4 (JIP4), and BLOC-one-related complex (BORC)) have been identified to mediate bidirectional lysosomal movement along microtubule tracks via kinesin and dynein.^18,19^ Among those, the amino acid-dependent regulation of lysosomal positioning by the interplay of Ragulator and BORC are among the best characterized. Intriguingly, this requires a dual role for the Ragulator complex, which either binds to the Rag GTPases, promoting mTORC1 signaling, or to BORC, regulating lysosomal motility, respectively.^20–22^ How this bifunctionality is regulated remains unknown to date. Regarding cholesterol, it has been known since the late 1990s^23^ that low levels promote perinuclear static lysosomes with increased ER contact sites. While the connection to mTORC1 signaling is increasingly well understood in this context,^8,16^ the mechanistic details of lysosomal positioning remain poorly characterized. First evidence demonstrated an important role of lysosomal transmembrane protein 55B (TMEM55B)-dependent recruitment of JIP4 in this context.^22^ A potential role for Ragulator/BORC, however, remains to be identified. Here, we characterized the cellular response to impaired lysosomal cholesterol homeostasis. We identified that mTOR phosphorylates the Ragulator complex member LAMTOR1 at serine 56 (S56) in response to altered lysosomal membrane cholesterol levels. We show that this cholesterol-specific phosphorylation event plays a critical role for mTORC1 signaling and lysosomal transport. Lamtor1 S56 phosphorylation displaces the Rag GTPases, and hence mTORC1 and enables binding of BORC, which results in perinuclear localization of lysosomes. Our data imply that LAMTOR1 S56 phosphorylation acts as a negative feedback mechanism, limiting mTORC1-dependent anabolic signaling in response to low lysosomal cholesterol levels.

## Results

### Inhibition of lysosomal cholesterol export impacts cellular phosphorylation dynamics and metabolite levels

Alterations of lysosomal cholesterol homeostasis affect subcellular lysosomal distribution.^24^ In accordance, we observed perinuclear clustering of lysosomes in mouse embryonic fibroblasts (MEF), HeLa cells, and human osteosarcoma (U2OS) cells concomitant with lysosomal cholesterol accumulation upon treatment with U18666A, an amphipathic steroid widely used to inhibit the lysosomal cholesterol transporter NPC1 (Figures 1A, 1B, S1 A-C).^25,26^ Importantly, unlike genetic models such as NPC1/2 KO cells, this system allows investigation of the acute effects of perturbations in lysosomal cholesterol handling. We used this setup to investigate cellular protein expression, phosphorylation-based signaling, and metabolism using a mass spectrometry (MS)-based multi-omics approach (Figure 1C). Lysosomes are low-abundant organelles (one cell typically contains a few hundred lysosomes),^27^ and lysosomal proteins have been estimated to comprise only ∼0.2% of cellular protein mass,^28^ resulting in reduced performance in whole cell proteomic studies.^29^ A frequently used approach to overcome this limitation is lysosome enrichment, for which superparamagnetic iron oxide nanoparticles (SPIONs) enable superior performance with respect to the selective recovery of terminal lysosomes.^30,31^ Hence, to cover both global and lysosome-centric processes in our proteomic datasets, we performed protein and phosphorylation level quantification from whole cell lysate (WCL) and lysosome-enriched fractions (LEF) following the stable isotope labeling by amino acids in cell culture (SILAC)^32^ approach (Table S1).

**Figure 1.**
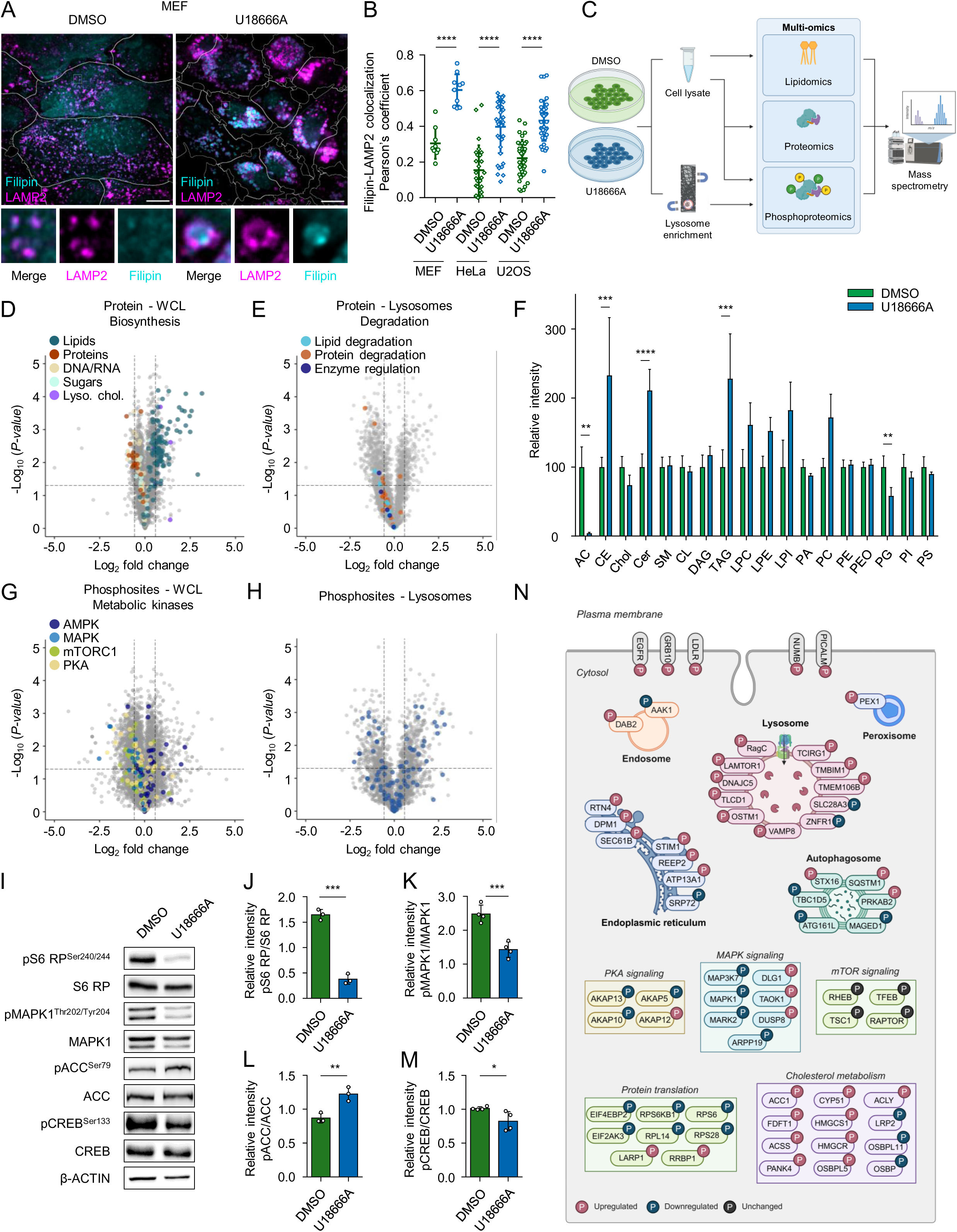
Multi OMICs analysis of impaired lysosomal cholesterol export identifies altered cellular response mechanisms. (A) U18666A induces lysosomal cholesterol accumulation. MEFs were treated with U18666A (3 µg/mL) or DMSO (0.03%) for 24 h followed by LAMP2 antibody and Filipin staining and confocal microscopy imaging. Magnified inset shown at the bottom. Scale bar: 10 µm. (B) Colocalization analysis of Filipin and LAMP2 in MEF, HeLa and U2OS cells using Fiji. The Pearson’s correlation coefficient between LAMP2 and Filipin is plotted on the y-axis for individual cells (n=9-20). Data represented are mean ± SD. Statistical significance was determined using one-way ANOVA followed by Sidak’s multiple comparisons test. **** p < 0.0001. (C) Experimental setup for multi-OMICs analysis of U18666A treated MEFs. Created with BioRender.com. (D) Lysosomal cholesterol buildup rewires biosynthetic pathways. Protein level changes in WCL, as determined by proteomic analysis of U18666A-treated versus DMSO-treated MEF cells. Differentially regulated proteins involved in lipid, protein, glyco, nucleic acid, and lysosomal cholesterol homeostasis (Lyso. chol.) identified by GSEA are highlighted. (E) Lysosomal cholesterol accumulation downregulates degradative pathways. Protein level changes in LEF, as determined by proteomic analysis of U18666A-treated versus DMSO-treated MEF cells. Differentially regulated proteins associated with protein/lipid degradation and enzyme regulation identified by GSEA are highlighted. (F) Targeted MS analysis of lipids in WCL of U18666A or DMSO-treated MEFs (n=5). AC, acylcarnitines; CE, cholesterol esters; Chol, cholines; Cer, ceramides; SM, sphingomyelins; CL, cardiolipins; DAG, diacylglycerols; TAG, triacylglycerols; LPC, lysophosphatidylcholines; LPE, lysophosphatidylethanolamines; LPI, lysophosphatidylinositols; PA, phosphatidic acids; PC, phosphatidylcholines; PE, phosphatidylethanolamines; PG, phosphatidylglycerols; PI, phosphatidylinositols; PS, phosphatidylserines. Data presented as mean ± SD. Statistical significance was determined by unpaired two-tailed t-test. ** p < 0.01, *** p < 0.001, and **** p < 0.0001. (G) Lysosomal cholesterol buildup alters phosphorylation-based signaling pathways. Protein phosphorylation levels in WCL, as determined by phosphoproteomic analysis of U18666A-treated versus DMSO-treated MEF cells. Differentially regulated phosphorylation sites of known substrates of the metabolic kinases AMP-activated protein kinase (AMPK), mitogen-activated protein kinase (MAPK), mechanistic target of rapamycin complex 1 (mTORC1), and protein kinase A (PKA) are highlighted (based on manual curation). (H) Regulation of lysosome-related protein phosphorylation levels. Protein phosphorylation levels in LEF, as determined by phosphoproteomic analysis of U18666A-treated versus DMSO-treated MEF cells. Differentially regulated phosphorylation sites of lysosomal or lysosome-associated proteins are highlighted based on a curated list from Hirn et al.^107^ (I) Activity of major metabolic kinases in response to lysosomal cholesterol buildup. MEF cells treated with U18666A or DMSO for 24 h, and cell lysates were probed with indicated antibodies. (J, K, L, M) Densitometric quantification of western blots from (I). Data are shown as mean ± SD (n=3-4 biological replicates, indicated by data points) of relative band intensities. Statistical significance was determined by unpaired two-tailed t-test. * p < 0.05, ** p < 0.01, *** p < 0.001, and **** p < 0.0001. (N) Comprehensive overview of differentially regulated phosphorylation sites in response to lysosomal cholesterol accumulation induced by U18666A in MEF cells. Created with BioRender.com.

In WCL and LEF, 210/172 and 75/507 proteins were up-/downregulated, respectively, following U18666A-induced lysosomal cholesterol accumulation. On a cellular level, mainly proteins involved in lipid biosynthesis were upregulated, while those related to the biosynthetic pathways of other basic cellular building blocks were either unregulated or reduced, including general protein biosynthesis, despite the presence of all other nutrients such as glucose or amino acids (cells were grown in full medium). In line with this, we observed members of both ribosomal subunits as strongest enriched categories in gene set enrichment analyses (GSEA) of downregulated proteins (Figure 1D, S1D, S1E, and Table S2). Further targeted investigation revealed coordinated upregulation of multiple key members of the cholesterol biosynthetic pathway (Figure S1F), in accordance with the previously demonstrated SREBP-mediated response to cellular cholesterol shortage as consequence of U18666A-based inhibition of lysosomal cholesterol efflux.^33^ Despite the strong accumulation of cholesterol in lysosomes (Figure 1A and 1B), and the expected impairment of lysosomal function,^34^ we did not detect upregulation of any lysosomal hydrolases, also such related to lipid hydrolysis (Figure 1E), suggesting no immediate activation of the transcription factor EB (TFEB) regulatory network^35^ under these conditions.

Concomitantly with the marked upregulation of proteins related to lipid biosynthesis (Figure 1D), MS-based lipidomic analyses identified an accumulation of various lipid species such as cholesteryl esters, diacylglycerols, triacylglycerols, and ceramides (Figure 1F, Table S3). The strong increase of cholesteryl esters and di-/triacylglycerol species (Figure S1G-S1I) was reminiscent of an increase in storage lipids, while phosphatidylethanol-amines and -inositols remained constant, indicating no radical changes in general membrane composition (Figure S1J and S1K). The accumulation of storage lipids, especially cholesteryl esters, could be due to a lack of their lysosomal degradation as consequence of U18666A treatment, as the resulting accumulation of free cholesterol in lysosomes (Figure 1A) was shown to result in a general impairment of lysosomal function. To differentiate between a malfunction of this catabolic pathway and the upregulation of cholesteryl ester formation as response to excess cholesterol biosynthesis,^36^ we investigated LEFs for cholesteryl ester accumulation by thin-layer chromatography (TLC). TLC analysis of WCL and LEFs following U18666A treatment identified that lysosomes predominantly accumulate free cholesterol, as also demonstrated by fillipin staining (Figure 1A) but not its esterified form (Figure S1L). Together with the observed protein regulatory patterns, this provides evidence for a strong upregulation of cholesterol biosynthesis and the acyl-CoA:cholesterol acyltransferase-mediated esterification of excess cholesterol produced by de novo synthesis. Additionally, we observed that U18666A-treated cells were largely depleted of acylcarnitines and carnitines (Figure 1F and S1M), which are crucial for shuttling fatty acids across the mitochondrial membrane to enable β-oxidation. As acetyl-CoA levels remained virtually unchanged between control cells and such treated with U18666A (Figure S1N), we interpreted this as a consequence of an increased flux of fatty acids into β-oxidation to provide acetyl-CoA for cholesterol de novo synthesis, as this process was shown previously to be decisive under lipid-starved conditions.^37,38^ Taken together, our data indicate a cholesterol shortage-dependent rerouting of cellular metabolism with reallocation of energetic resources to cholesterol biosynthesis.

It is well-established that the activity of cellular metabolic pathways, and hence metabolic flux, is mediated by several kinases, including AMP-activated protein kinase (AMPK), mTORC1, protein kinase A (PKA), and MAPK, which integrate information from both intracellular metabolite levels and extracellular signals.^10,39^ How lysosomes contribute to this network of cellular signaling, particularly in response to cholesterol, has not been fully characterized. To monitor phosphorylation dynamics in relation to cellular metabolism and lysosome-related proteins as a consequence of lysosomal cholesterol accumulation, we performed phosphoproteomics analysis of WCL and LEF following U18666A treatment (Table S4). In WCL and LEF, 325/814 and 351/330 phosphorylation sites showed increased/decreased levels, respectively (Figure 1G and 1H). This included downregulated phosphorylation of substrates for mTORC1 (S6 ribosomal protein, S6 RP), PKA (cAMP response element binding protein, CREB), and MAPK (MAPK1), alongside upregulation of phosphorylation of the AMPK substrate acetyl-CoA carboxylase (ACC), reflecting its well-characterized opposing role to mTORC1. We corroborated these findings by western blot analyses using phosphosite-specific antibodies for thoroughly validated substrates of individual kinases (Figure 1I-M). Importantly, the observed phosphorylation dynamics also correlated with patterns observed in protein level data, with a strong upregulation of protein phosphorylation of members of the lipid biosynthetic pathway (Figure S1O and S1P). Gene ontology (GO) analysis further revealed regulation of phosphorylation sites related to intracellular vesicles, transport complexes, and membrane dynamics (Figure S1Q and S1R). Importantly, several of these sites were localized at lysosome-related proteins, for which we identified 165 phosphorylation sites in total. This included 55 regulated sites on lysosomal protein domains exposed to the cytosol, representing possible regulators of lysosomal motility and positioning, as well as sites on other subcellular compartments (Figure 1H and 1N).

Taken together, our multi-omics analysis of lysosomal cholesterol egress blockage through inhibition of NPC1 by U18666A reveals widespread changes in protein abundance and phosphorylation, especially for those related to cholesterol metabolism, accompanied by reduced levels of proteins related to other biosynthetic pathways. Importantly, we identify several phosphorylation sites on lysosomal surface-exposed proteins that are co-regulated with the main actors in cholesterol biosynthesis, implying putative common upstream regulatory processes.

### LAMTOR1 S56 phosphorylation displaces Rag GTPases from the Ragulator complex

Based on these data, we hypothesized that differential phosphorylation of lysosomal surface proteins could be a critical factor in the regulation of lysosomal positioning in response to altered cholesterol homeostasis, as similar mechanisms are known for mitochondria and peroxisomes.^40,41^ Indeed, we identified several regulated phosphorylation sites on lysosomal surface proteins/complexes which were previously connected with lysosomal motility,^24^ vesicle fusion,^42^ or cholesterol exchange.^43^ As it was shown that lysosomal transport is dependent on the assembly of protein complexes, we investigated phosphorylation-dependent complex assembly. As readout, we used protein half-life determination, as it was previously shown that complex subunits are stabilized upon proper incorporation.^44,45^ We selected four differentially regulated lysosomal phosphorylation sites from our dataset (LAMTOR1-S56, RagC-S381, STARD3NL-S39, and VAMP8-T54) for further investigation, and generated phosphomimetic/-resistant (S/T to E/A) versions, respectively. We assessed the stability of wildtype (WT), E, and A versions by transfecting human embryonic kidney 293 cells (HEK293) cells followed by cycloheximide treatment^46,47^ and western blot detection (Figure 2A). While the phosphomimetic/-resistant versions of RagC-S381 and STARD3NL-S39 did not affect the stability of the respective proteins (Figure S2A and S2B), VAMP8 showed increased stability for both T54E and T54A (Figure S2C). Notably, this amino acid residue has previously been reported to regulate autophagosome-lysosome fusion.^42^ Also, LAMTOR1 stability was markedly increased for the S56A mutant compared to both WT and S56E (Figure 2B).

**Figure 2.**
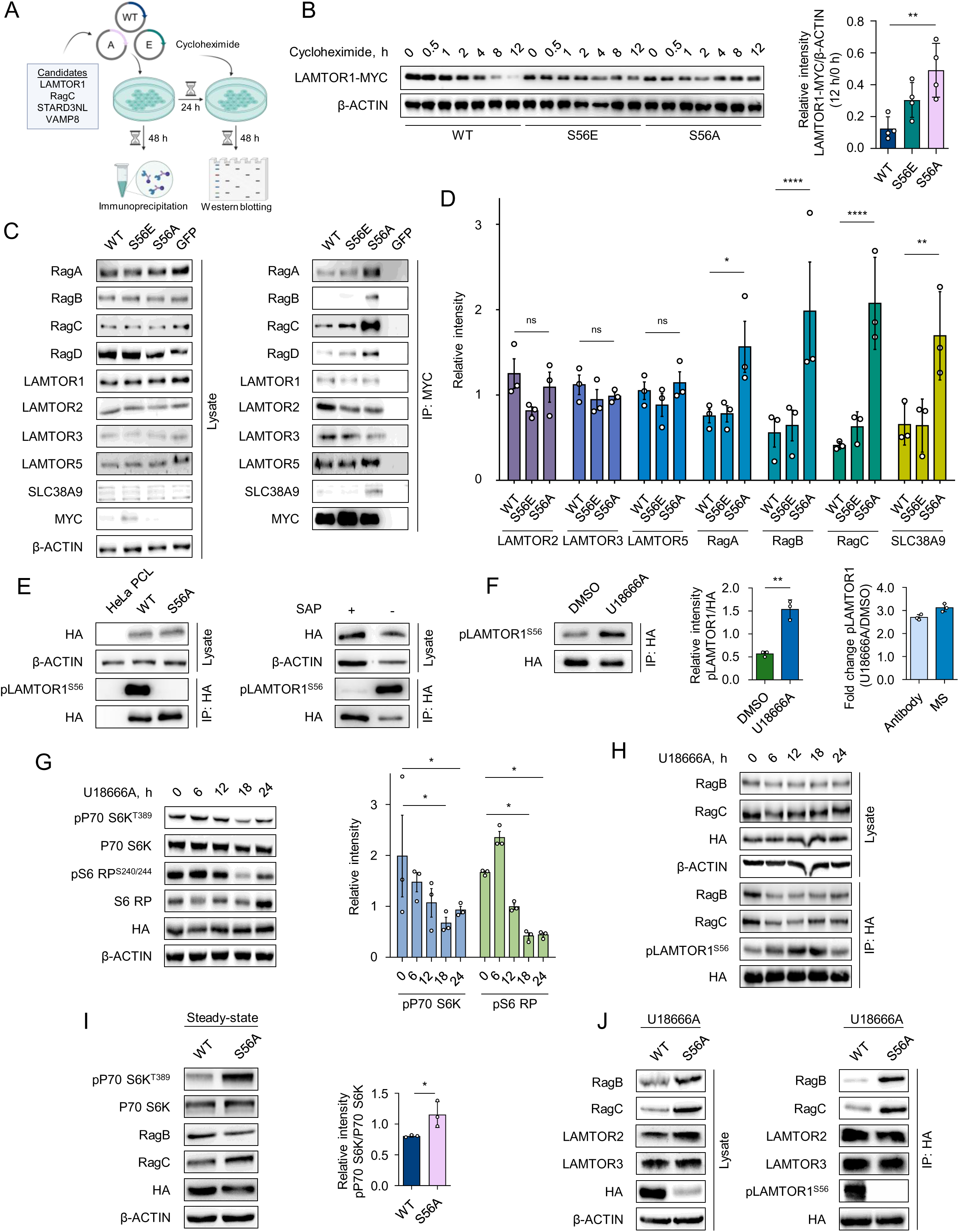
LAMTOR1 phosphorylation affects Ragulator-Rag GTPases interaction. (A) Experimental setup of (B) and (C). Created with BioRender.com. (B) Phosphorylation status of LAMTOR1 S56 affects the protein’s stability. HEK293 cells were transfected with MYC-tagged LAMTOR1 WT, S56E, or S56A followed by incubation with cycloheximide for the specified time. Left: Cell lysates were probed with the indicated antibodies. Right: Densitometric quantification of MYC-tagged LAMTOR1 WT, S56E, and S56A protein levels of the 12 h time point, normalized to 0 h. Data shown are mean ± SD (n=4 biological replicates, indicated by data points) of relative band intensities. Statistical significance was determined using one-way ANOVA followed by Dunnett’s multiple comparisons test. ** p < 0.01. (C) LAMTOR1 S56 phosphorylation displaces Rag GTPases and SLC38A9 from the Ragulator complex. HEK293 cells were transected with MYC-tagged LAMTOR1 WT, S56E, S56A, or GFP followed by MYC-IP. IP eluates and cell lysates were immunoblotted and probed with indicated antibodies. (D) Densitometric quantification of immunoblots from (C). Shown are mean values ± SD (n=3 biological replicates, indicated by data points) of relative band intensities. Statistical significance was determined using one-way ANOVA followed by Dunnett’s multiple comparisons test. * p < 0.05, ** p < 0.01, and **** p < 0.0001. (E) Validation of LAMTOR1 S56 phosphorylation-specific antibody. Left: HeLa parental cell line (PCL), HA-tagged LAMTOR1 WT, or S56A cell lysates were used for HA-IP. Right: Cell lysates from HA-tagged LAMTOR1 WT were treated with shrimp alkaline phosphatase followed by HA-IP. In both cases, IP eluate fractions were analyzed for phosphorylation status of LAMTOR1 S56 and total protein abundance. (F) Validation of LAMTOR1 S56 phosphorylation levels determined by phosphosite-specific antibody versus phosphoproteomic analysis. Left: HA-tagged LAMTOR1 WT cells were treated with U18666A (3 µg/mL) or DMSO (0.03%) as control for 24 h followed by HA-IP. IP eluates were analyzed for phosphorylation status of LAMTOR1 S56 and total protein abundance. Middle: Densitometric quantification of immunoblots, shown are mean values ± SD (n=3 biological replicates, indicated by data points) of relative band intensities. Statistical significance was determined using unpaired two-tailed t-test. ** p < 0.01. Right: Comparison of fold changes in LAMTOR1 S56 phosphorylation levels determined by immunoblotting and mass spectrometric analysis. (G) Time-dependency of U18666A on canonical mTORC1 activity. Left: HA-tagged LAMTOR1 WT cells were treated with U18666A (3 µg/mL) for different periods, as indicated. Cell lysates were probed with specified antibodies. Right: Densitometric quantification of immunoblots, shown are mean values ± SEM (n=3 biological replicates, indicated by data points) of relative band intensities. Statistical significance was determined using two-way ANOVA followed by Dunnett’s multiple comparisons test. * p < 0.05. (H) Time-dependency of U18666A on Rag GTPase-Ragulator interaction. HA-tagged LAMTOR1 WT cells were treated with U18666A (3 µg/mL) for different periods, as indicated and cell lysates were subjected to HA-IP. IP eluates and cell lysates were probed with antibodies as indicated. (I) Constitutive activation of canonical mTORC1 in LAMTOR1 S56A. Left: Cell lysates from HA-tagged LAMTOR1 WT or S56A were probed with antibodies as indicated. Right: Densitometric quantification of phosphorylation levels of P70 S6K; shown are mean values ± SD (n=3 biological replicates, indicated by data points) of relative band intensities. Statistical significance was determined using unpaired two-tailed t-test. * p < 0.05. (J) HA-tagged LAMTOR1 WT or S56A cells were treated with U18666A (3 µg/mL) for 24 h followed by HA-IP. IP eluates and cell lysates were subjected to immunoblotting using the indicated antibodies. Phosphorylation levels of all proteins detected by phospho-specific antibodies in immunoblots were quantified and normalized to their corresponding total protein levels.

LAMTOR1 forms with LAMTOR2/3/4/5 the heteropentameric Ragulator complex, which is of importance for both lysosomal signaling and positioning, through interaction with members of mTORC1 or BORC, respectively.^20,21^ To facilitate mTORC1 signaling, Ragulator interacts with the vacuolar-type ATPase (V-ATPase or H+-ATPase) complex, the Rag GTPase heterodimers RagA/RagB and RagC/RagD, as well as the amino acid transporter SLC38A9, which mediates nutrient-dependent mTORC1 activation at the lysosomal surface.^48,49^ Therefore, we initially tested whether LAMTOR1 pS56 affects assembly of the Ragulator complex and/or the mTORC1 nutrient sensing complex in HEK293 cells through co-immunoprecipitation (co-IP) of MYC-tagged LAMTOR1 WT, S56E, and S56A followed by immunoblot detection of the other complex members (Figure 2C). While LAMTOR1 S56A did not affect assembly of the Ragulator complex, it increased the affinity for Rag GTPases and SLC38A9 compared to WT, the S56E mutant had no effect (Figure 2D). These findings suggest that LAMTOR1 pS56 could disrupt the interaction between Ragulator and the Rag GTPases/SLC38A9, facilitating deactivation of mTORC1, which we also detected under conditions of increased LAMTOR1 pS56 levels (Figure 1J). After manual inspection of MS/MS spectra to confirm LAMTOR1 pS56 identity (excluding S55 phosphorylation) (Figure S2D), and we verification of its conservation across species (Figure S2E), we generated LAMTOR1 knock down HeLa cells^50^ stably expressing HA-tagged LAMTOR1 S56 WT, E, and A (Figure S2F). In these cells, we confirmed our previous results for constant/differential binding of LAMTOR5 and RagA/C, respectively (Figure S2G). Importantly, we further generated a LAMTOR1 pS56-specific rabbit antibody (Figure S2H and S2I) and validated it through differential detection of LAMTOR1 S56 WT and S56A, as well as *in vitro* dephosphorylation of LAMTOR1 S56WT (Figure 2E). Finally, we showed that the phospho-specific antibody was able to quantify changes in LAMTOR1 pS56 levels upon U18666A treatment and that the observed changes in LAMTOR1 S56 phosphorylation levels correlated with our MS data (Figure 2F).

To investigate how lysosomal cholesterol accumulation affects mTORC1 signaling over time, and how it correlates with LAMTOR1 S56 phosphorylation and Rag GTPase binding, we performed a 24 h U18666A time course experiment in LAMTOR1 S56WT cells. We monitored mTORC1 activity through phosphorylation of its canonical substrates 70-kilodalton ribosomal protein S6 kinase (P70 S6K) and S6 RP, and examined the relationship between LAMTOR1 pS56 and Rag GTPase binding through co-IP experiments (Figure 2G and 2H). This allowed us to determine whether LAMTOR1 S56 phosphorylation correlates with changes in mTORC1 activity and Rag GTPase recruitment during cholesterol-induced lysosomal stress. In accordance with our steady state co-IP results for different LAMTOR1 versions (Figure 2D), we identified an inverse correlation between LAMTOR S56 phosphorylation (increase) and binding of RagB/RagC, and hence mTORC1 activity (decrease). These data provide further evidence that LAMTOR1 S56 phosphorylation could be responsible for mTORC1 deactivation through displacement of Rag GTPases, placing this phosphorylation event upstream of mTORC1. In accordance, we identified higher constitutive mTORC1 activity in LAMTOR1 S56A cells under steady state conditions (Figure 2I). Consistently, upon U18666A treatment, we observed higher RagB/RagC affinity at constant Ragulator composition (Figure 2J), correlating with constitutive activation of mTORC1.

These data indicate that LAMTOR1 S56 phosphorylation is responsible, and sufficient, to deactivate mTORC1 through displacement of Rag GTPases in a cholesterol-dependent manner, assigning a central regulatory function to this residue. Lending further credibility to this hypothesis, LAMTOR1 S56 is located in a region of LAMTOR1 which is unstructured in its free state and adapts an alpha helical conformation only upon interaction with RagC.^50^ Furthermore, deletion of this protein region (AA 51-60) resulted in deactivation of mTORC1 while removal of the remaining LAMTOR1 N-terminus (AA 1-40) had no effect (Figure S2J).^51^

### LAMTOR1 pS56 regulates BORC-dependent lysosomal motility

In addition to its central role for mTORC1 signaling, Ragulator was shown to regulate lysosomal positioning through interaction with the hetero-octameric BORC complex.^21,52^ Under amino acid sufficiency, Ragulator engages in mutually exclusive interactions with either the Rag GTPases or BORC, giving rise to functionally distinct Ragulator complexes on the lysosomal membrane.^53^ Rag GTPase-bound Ragulator results in free BORC, which then interacts with the small GTPase ARL8B and the kinesin KIF5B, catalyzing the anterograde movement of lysosomes along microtubule tracks. Importantly, as Ragulator and the Rag GTPases are much more abundant than BORC,^28^ both populations (Rag GTPase- and BORC-associated Ragulator) co-exist on the lysosomal surface. It has been reported, that under amino acid insufficiency, Rag GTPases remain also associated with a pool of Ragulator in a stable but inactive configuration, promoting mTORC1 inhibition.^15,49^ This presents a scenario, where Ragulator is occupied with Rag GTPases both with and without amino acids. How it is possible under these conditions to displace the Rag GTPases, enabling binding of BORC to Ragulator, has not been identified to date. We hypothesized that LAMTOR1 S56 phosphorylation may be able to play a decisive role in this process.

Along this line, upon phosphorylation of LAMTOR1 at S56 and the concomitant displacement of Rag GTPases from Ragulator (Figure 2), we observed perinuclear clustering of lysosomes. This matches previous results showing that LAMTOR1 knockdown or knockout (KO) results in more static and peripheral lysosomes.^21^ To follow up on this, we visualized lysosomal distribution using lysosome-associated membrane glycoprotein 2 (LAMP2) staining in cells expressing HA-tagged LAMTOR1 WT and S56A, respectively. In line with our hypothesis, we identified a significant shift towards more peripheral lysosomal localization for cells expressing LAMTOR1 S56A relative to LAMTOR1 WT, including pronounced plasma membrane-localized lysosomal clusters (Figure 3A). In these cells, we performed siRNA-based knockdown of BORC subunit 5 (BORCS5) and ARL8B to investigate BORC’s role. After validation of siRNA knockdown efficiency through western blot-based detection of BORCS5/ARL8B protein levels (Figure S3A), we visualized lysosomal distribution through LAMP2 immunostaining. In these cells, it was possible to revert the LAMTOR1 S56A phenotype through knockdown of either BORCS5 or ARL8B (Figure 3B), confirming that the observed lysosomal peripheral localization (Figure 3A) was dependent on the lack of BORC-to-Ragulator-interaction, most likely as a consequence of LAMTOR S56A’s increased affinity to the Rag GTPases (Figure 2D).

**Figure 3.**
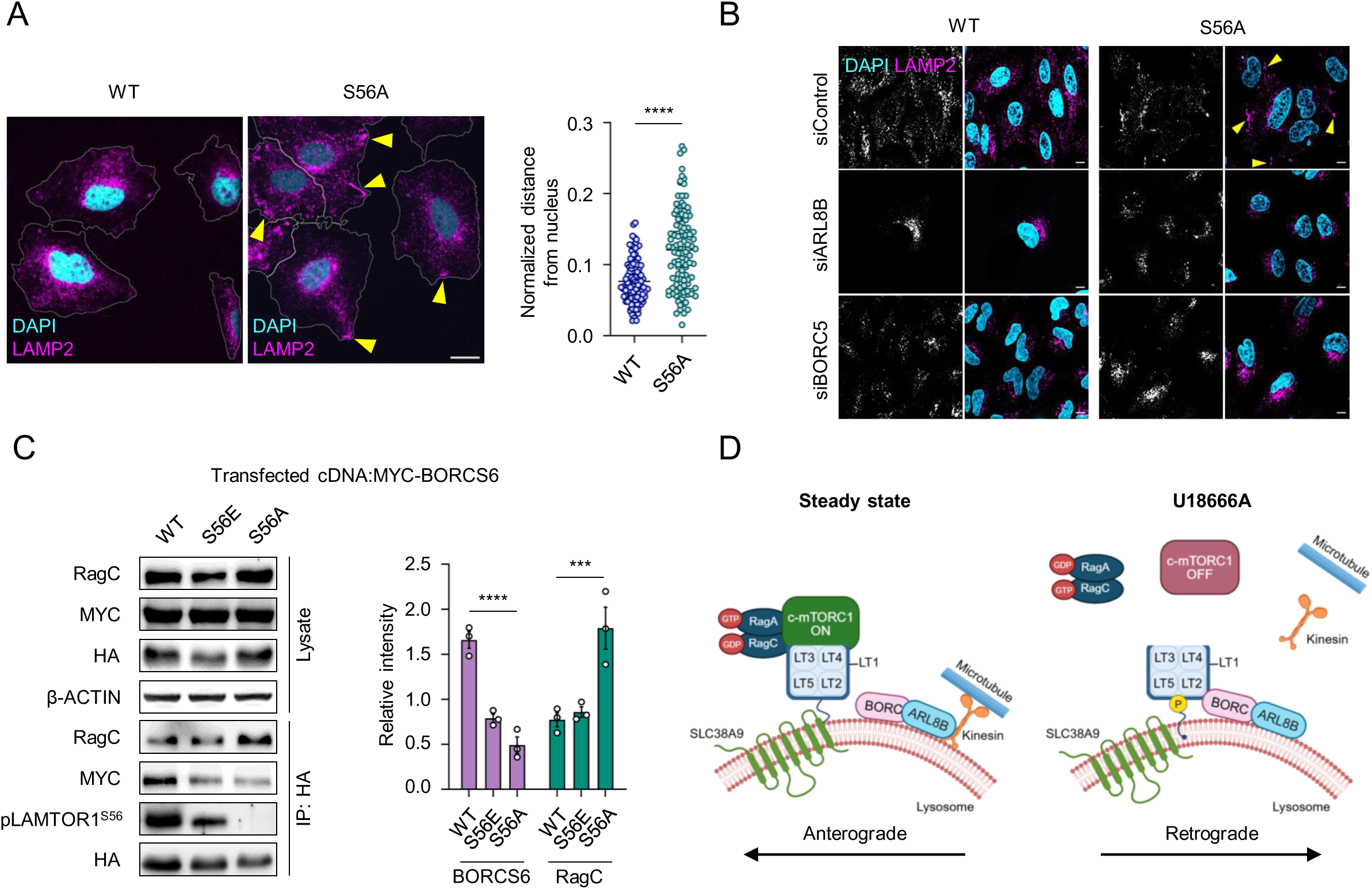
LAMTOR1 pS56 promotes retrograde lysosomal movement. (A) Increased peripheral clusters of lysosomes in LAMTOR1 S56A cells. HA-tagged LAMTOR1 WT or S56A cells were stained using anti LAMP2 antibodies and imaged by confocal microscopy. Arrows mark peripheral clusters of LAMP2-stained vesicles (lysosomes). Scale bar: 10 µM. Plotted are normalized distances of lysosomes relative to the nucleus for individual cells (n=116). Mean ± SD is indicated. Statistical significance was determined using unpaired two-tailed t-test. **** p < 0.0001. (B) LAMTOR1 S56-dependent regulation of lysosomal positioning depends on BORC. HA-tagged LAMTOR1 WT, or S56 cells were transfected with scrambled-, ARL8B- or BORCS5-siRNAs, stained using anti LAMP2 antibodies and imaged using fluorescence microscopy. (C) LAMTOR1 S56 phosphorylation regulates Ragulator-Rag GTPases/BORC binding. HA-tagged LAMTOR1 WT, S56E and S56A cells were transfected with MYC-BORCS6 cDNA followed by HA-IP. Left: Cell lysates and IP eluates were probed with antibodies, as indicated. Right: Densitometric quantification of BORCS6 (MYC) or RagC levels; shown are mean values ± SEM (n=3 biological replicates, indicated by data points) of relative band intensities. Statistical significance was determined using two-way ANOVA followed by Sidak’s multiple comparisons test. *** p < 0.001, **** p < 0.0001. (D) Influence of LAMTOR1 S56 phosphorylation on Ragulator/Rag GTPase and Ragulator/BORC binding. LAMTOR1 S56 phosphorylation displaces Rag GTPases, enabling binding of BORC to Ragulator and mediating lysosomal retrograde movement.

These data imply that the inverse binding relationship between BORC/Rag GTPases to individual Ragulator complexes could be regulated through phosphorylation of LAMTOR1 S56. To gain more insights into such a mechanism, we transfected HeLa cells expressing HA-tagged LAMTOR1 WT, S56E, and S56A with MYC-tagged BORC subunit 6 (BORCS6) and performed co-IP experiments. In line with our hypothesis, BORCS6 exhibited reduced affinity to LAMTOR1 S56A relative to S56E and WT (representing the phosphorylated protein) in HA-LAMTOR1-IPs, presenting with a S56 phosphorylation-dependent inverse correlation with respect to RagC (Figure 3C). Furthermore, we performed the inverse experiment with MYC-BORCS6 co-IPs from these cells, confirming the effect of LAMTOR1 S56 (Figure S3B). Finally, as these experiments were performed in overexpression systems due to the insufficient sensitivity of available antibodies to detect endogenous BORC, we investigated our proteomic dataset of LEFs (Table S1). Indeed, in line with the co-IP data, we identified a significant downregulation of ARL8B levels upon treatment with U18666A, indicative of Ragulator-bound, and hence inactive, BORC (Figure S3C).

These findings, together with the co-IP results of Rag GTPase binding to individual LAMTOR1 S56 versions, support a model in which phosphorylation of LAMTOR1 S56 acts as a context-dependent molecular switch, modulating the interaction of Ragulator with either the Rag GTPases or BORC (Figure 3D).

### Lysosomal membrane cholesterol levels regulate LAMTOR1 S56 phosphorylation

Previous studies have described changes in Ragulator-BORC interactions concurrently with mTORC1 inactivation upon amino acid starvation. Notably, it was demonstrated that the Rag GTPases are usually not fully displaced from Ragulator in response to amino acid starvation, but rather alter their residence time and GDP/GTP-dependent configuration, affecting mTORC1 acitivity^15^ and, therefore, possibly BORC binding. In our study, we utilized U18666A, an inhibitor of the widely recognized primary lysosomal cholesterol exporter NPC1 which incorporates intra-lysosomal NPC2-bound cholesterol into the lysosomal limiting membrane,^5^ triggering lysosomal cholesterol accumulation (Figure 1A). For several pathological conditions, such as Niemann Pick Disease Type C, it was demonstrated that lysosomal cholesterol accumulation results in secondary effects due to the overall impairment of lysosomal function.^54^ This raises the possibility that other stimuli than lysosomal cholesterol accumulation trigger LAMTOR1 S56 phosphorylation. To explore different input signals, we perturbed amino acid availability, total cellular cholesterol levels, lysosomal cholesterol flux, ER-lysosome-cholesterol exchange, and phosphatidylinositol 4-phosphate (PI(4)P) levels (Figure 4A).

**Figure 4.**
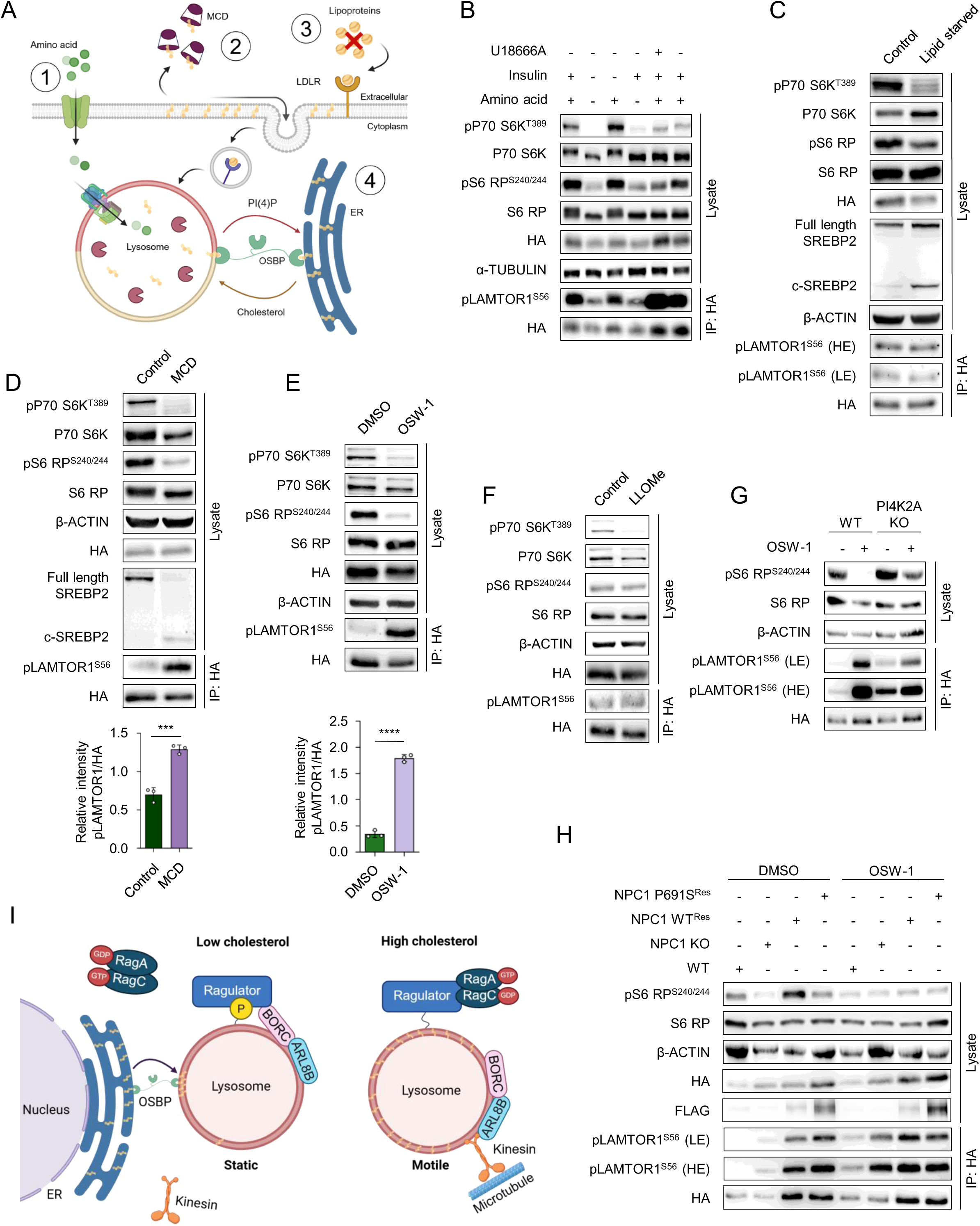
Loss of lysosomal limiting membrane cholesterol triggers LAMTOR1 pS56. (A) Model depicting the effect of 1) amino acid starvation, 2) MCD 3) LPDS and 4) OSW-1. Created with BioRender.com. (B) Amino acid starvation does not trigger LAMTOR1 S56 phosphorylation. HA-tagged LAMTOR1 WT cells were treated with full media (containing amino acids) with or without U18666A for 24 h (3 µg/mL) or starved for amino acids for 5 h, followed either by refeeding with amino acids for 30 min or stimulation with insulin for 10 min. IP eluates and cell lysates were probed with antibodies as indicated. (C) Reduction of lysosomal cholesterol flux through the endocytic pathway does not affect LAMTOR1 S56 phosphorylation. HA-tagged LAMTOR1 WT cells were lipid-starved by incubating in medium containing 3% LPDS for 17 h. IP eluates and cell lysates were probed with antibodies as indicated (D) Depletion of membrane cholesterol levels using MCD triggers LAMTOR1 s56 phosphorylation. HA-tagged LAMTOR1 WT cells were treated with 0.5% MCD for 2 h in combination with 3% LPDS. IP eluates and cell lysates were probed with antibodies as indicated. Densitometric quantification of band intensities of LAMTOR1 S56 phosphorylation was performed and data shown are mean ± SD (n=3 biological replicates, indicated by data points). Statistical significance was determined using unpaired two-tailed t-test. *** p < 0.001. (E) ER-lysosome cholesterol transport inhibition induces LAMTOR1 S56 phosphorylation. HA-tagged LAMTOR1 WT cells were treated with DMSO or 200 nM of OSW-1 for 2 h. IP eluates and cell lysates were probed with antibodies as indicated. Densitometric quantification of band intensities of LAMTOR1 S56 phosphorylation was performed, shown are mean values ± SD (n=3 biological replicates, indicated by data points). Statistical significance was determined using unpaired two-tailed t-test. **** p < 0.0001. (F) LLOMe-induced lysosomal PI(4)P accumulation does not trigger LAMTOR1 S56 phosphorylation. HA-tagged LAMTOR1 WT cells were treated with 1 mM LLOMe for 30 min. IP eluates and cell lysates were probed with antibodies as indicated. (G) PI(4)P synthesis does not affect LAMTOR1 phosphorylation. U2OS WT or U2OS PI4K2A KO cells expressing HA-tagged LAMTOR1 WT were treated with DMSO or 200 nM OSW-1 for 2 h. IP eluates and cell lysates were probed with antibodies as indicated. (H) Depletion of lysosomal membrane cholesterol regulates LAMTOR1 S56 phosphorylation. HEK293 cells expressing HA-tagged LAMTOR1 WT along with the other indicated genotypes were treated with DMSO or 200 nM OSW-1 for 2 h. IP eluates and cell lysates were probed with antibodies as indicated. (I) Model for regulation of lysosomal membrane cholesterol-dependent lysosomal retrograde positioning via LAMTOR1 S56 phosphorylation. Depletion of lysosomal membrane cholesterol triggers LAMTOR1 S56 phosphorylation regulating the movement of lysosomes toward perinuclear region. Created with Biorender.com. For all experiments, cell lysates were used to assess canonical mTORC1 activity by examining the phosphorylation status of its downstream targets, P70 S6K and/or S6 RP, as indicated, while IP eluates from the same samples were used to monitor LAMTOR1 S56 phosphorylation.

Initially, we determined the relationship of LAMTOR1 S56 phosphorylation and canonical mTORC1 activation through amino acid starvation/refeeding and insulin stimulation (Figure 4B). Under amino acid starvation, which resembles canonical mTORC1 inactivation, we observed reduced LAMTOR1 pS56 phosphorylation levels, suggesting that it is unlikely to be driven by canonical mTORC1. A possible explanation for the observed reduction of LAMTOR1 S56 phosphorylation is that amino acid starvation results in altered affinity of Rag GTPases,^20,21^ which suffices to free the Ragulator complex for binding of BORC irrespective of LAMTOR1 S56 phosphorylation. Further substantiating this finding, insulin stimulation was insufficient to recover LAMTOR1 S56 phosphorylation levels (Figure 4B). It was shown previously, that lysosomal cholesterol levels, sensed through NPC1/SLC38A9 and LYCHOS, are decisive for mTORC1 activity.^11,13^ We investigated involvement of these sensory mechanisms by treating cells with lipid-depleted serum (LPDS),^55^ reducing cholesterol flux through the endolysosomal pathway. Increased SREBP2 proteolytic processing and concomitant downregulation of mTORC1 confirmed reduced lysosomal cholesterol delivery^55,56^ (Figure 4C). Also, in these experiments, we did not observe upregulation of LAMTOR1 S56 phosphorylation, providing further evidence that the lysosomal surface localized mTORC1-related nutrient sensing machinery is not triggering LAMTOR1 S56 phosphorylation, also not in response to altered levels of lysosomal cholesterol flow.

To not only affect endolysosomal cholesterol flux, but the cholesterol content of cellular membranes, we treated cells with methyl-β-cyclodextrin (MCD),^11^ a cyclic heptasaccharide which depletes cholesterol from the plasma membrane, affecting overall cellular cholesterol homeostasis. Consistent with the literature, MCD treatment reduced mTORC1 activity, activated SREBP2,^57^ and, importantly, significantly increased LAMTOR1 S56 phosphorylation (Figure 4D), implying that it is triggered in response to reduced membrane cholesterol levels rather than endolysosomal cholesterol flow. We further performed LAMTOR1 co-IP experiments of MCD-treated cells, observing the same reduced affinity of Rag GTPases at constant Ragulator complex composition as for inhibition of lysosomal cholesterol egress by U18666A (Figure S4A). Strikingly, in LAMTOR1 S56A-expressing cells MCD treatment failed to displace the Rag GTPases, and hence to deactivate mTORC1, implying a crucial role of LAMTOR1 S56 phosphorylation for the deactivation of canonical mTORC1 in response to membrane cholesterol depletion (Figure S4A). A common feature of MCD and U18666A is the reduction of lysosomal membrane cholesterol levels, either through reduced ER to lysosome transport due to lower ER cholesterol levels (MCD) or reduced lysosomal lumen to membrane shuttling (U18666A). We, therefore, hypothesized that lysosomal membrane cholesterol content, and its maintenance through ER-derived cholesterol, could affect LAMTOR1 S56 phosphorylation. To address this, we inhibited ER to lysosome cholesterol transport with the OSBP-specific inhibitor OSW-1.^8,58^ In line with previous studies,^16^ blockage of ER cholesterol shuttling with OSW-1 reduced mTORC1 activity and, strikingly, resulted in a strong induction of LAMTOR1 S56 phosphorylation (Figure 4E).

Mechanistically, OSW-1 inhibits the OSBP-mediated exchange of ER cholesterol for lysosomal PI(4)P at ER-lysosome contact sites. This exchange is driven by SAC1, an ER-resident PI(4)P-phosphatase, which generates a PI(4)P gradient driving cholesterol transport towards lysosomes. Importantly, loss of SAC1, and the resulting disruption of ER-lysosome cholesterol transport, has been shown to affect lysosomal motility, and elevated PI(4)P levels in perinuclear lysosomes have been linked to mTORC1 activity.^16^ OSW-1 based inhibition of ER to lysosome cholesterol exchange could, therefore, act on LAMTOR1 S56 through i) reduction of ER-derived cholesterol on the lysosomal limiting membrane or ii) increase of lysosomal PI(4)P levels.^16,59^ To discriminate between these scenarios, we treated cells with L-leucyl-L-leucine methyl ester hydrobromide (LLOMe), which causes lysosomal damage that triggers lysosomal PI(4)P generation and, consequentially, increased ER to lysosome cholesterol flux via ORP1L.^60^ While LLOMe, consistent with previous results, reduced mTORC1 activity,^16^ it did not alter LAMTOR1 pS56 levels (Figure 4F). Furthermore, we detected increased levels of LAMTOR1 S56 phosphorylation in OSW-1 treated U2OS Phosphatidylinositol 4-kinase type 2-alpha (PI4K2A) KO cells (Figure S4B) demonstrating that regulation of LAMTOR1 S56 phosphorylation is independent of PI(4)P (Figure 4G). These data further confirm that reduced mTORC1 activity alone is not sufficient to trigger LAMTOR1 S56 phosphorylation, that it is regulated independent of lysosomal PI(4)P levels, and that it exclusively responds to a reduction of lysosomal limiting membrane cholesterol levels.

Finally, we confirmed the decisive role of ER to lysosome cholesterol transport for LAMTOR1 S56 phosphorylation through utilization of NPC1 KO cells. It was shown previously, that NPC1 KO leads to increased ER-lysosome contact sites, facilitating a continuous delivery of ER-derived cholesterol to lysosomes as compensation for the lack of LDL-derived cholesterol supply.^8^ Given the high ER to lysosome flux of endogenously synthesized cholesterol under steady state conditions in these cells, we hypothesized that they would have a reduced requirement for LAMTOR1 S56 phosphorylation upon membrane cholesterol depletion. Using CRISPR-Cas9, we generated HEK293 NPC1 KO cells (Figure S4C) expressing HA-tagged LAMTOR1 WT and treated them with MCD. In line with our hypothesis, this treatment did not induce any changes in LAMTOR1 S56 phosphorylation levels (Figure S4D). Strikingly, when we treated these cells with OSW-1, blocking their major source of cholesterol for the lysosomal limiting membrane, we observed a strong induction of LAMTOR1 S56 phosphorylation (Figure 4H). To further address if the effect of NPC1 KO was indeed due to altered lysosomal membrane cholesterol levels, or downstream of NPC1-mediated cholesterol sensing, and hence mTORC1 signaling, we reconstituted NPC1 KO cells with WT NPC1 and the sterol sensing mutant NPC1 P691S, which was shown previously to be unable to transmit cholesterol levels to mTORC1. Cells expressing NPC1 P691S showed no difference with respect to LAMTOR1 pS56 levels upon OSW-1 treatment compared to such reconstituted with NPC1 WT (or NPC1 KO cells), further lending the hypothesis that alterations of lysosomal membrane cholesterol levels in response to ER-to-lysosome cholesterol exchange present the decisive factor (Figure 4H).

Taken together, these results suggest that phosphorylation of LAMTOR1 S56 is exclusively induced in response to a shortage of lysosomal membrane cholesterol. This results in displacement of the Rag GTPases from Ragulator, which in turn binds to the BORC, resulting in static perinuclear lysosomes which can efficiently engage in ER contact sites, enabling replenishment of lysosome membrane cholesterol levels via OSBPs (Figure 4I).

### mTOR phosphorylates LAMTOR1 at serine 56

Our data imply that LAMTOR1 S56 phosphorylation is part of a regulatory network functioning independent of (canonical) mTORC1 signaling which is, to our knowledge, currently the only signaling pathway known to be involved in lysosomal cholesterol signaling.^8,11,13^ We, therefore, set out to further investigate the regulation of LAMTOR1 S56 phosphorylation. Initially, based on the regulatory patterns we identified in our MEF U18666A experiments (Figure 1G and 1I to 1M), we tested the effect of major metabolic signaling kinases on LAMTOR1 S56 phosphorylation.

AMPK is well-known for its activation under conditions of low cellular energy levels and was the only kinase for which we found increased activity upon perturbation of lysosomal cholesterol homeostasis (Figure 1H and 1L). Also, its activity was shown to be negatively correlated with mTORC1,^61^ and it can be activated at the lysosomal membrane in a Ragulator-dependent fashion.^62^ We, therefore, stimulated/inhibited AMPK using the small molecules A769662^63^ or dorsomorphin,^64^ respectively, and monitored phosphorylation of its well-established downstream substrate Acetyl-CoA-Carboxylase (ACC).^65^ While AMPK activation did not affect LAMTOR1 S56 phosphorylation, surprisingly, its inhibition resulted in increased levels (Figure 5A), indicating an inverse relationship. We further investigated PKA, whose activity also was shown to be negatively correlated with mTORC1,^66^ trough activation/inhibition using forskolin/IBMX^66^ or H89,^67^ respectively. As a readout for PKA activity, we monitored phosphorylation of its direct substrate cAMP-Response Element-Binding Protein (CREB).^68^ Also in these experiments, we found an inverse correlation of PKA activity and LAMTOR1 S56 phosphorylation (Figure 5B). Finally, we investigated MAPK1, which was demonstrated to directly interact with LAMTOR1,^69^ via activation/inhibition by Epidermal Growth Factor (EGF) and U0126,^70^ respectively (Figure 5C). In these experiments, we did not detect any variations in LAMTOR1 S56 phosphorylation.

**Figure 5.**
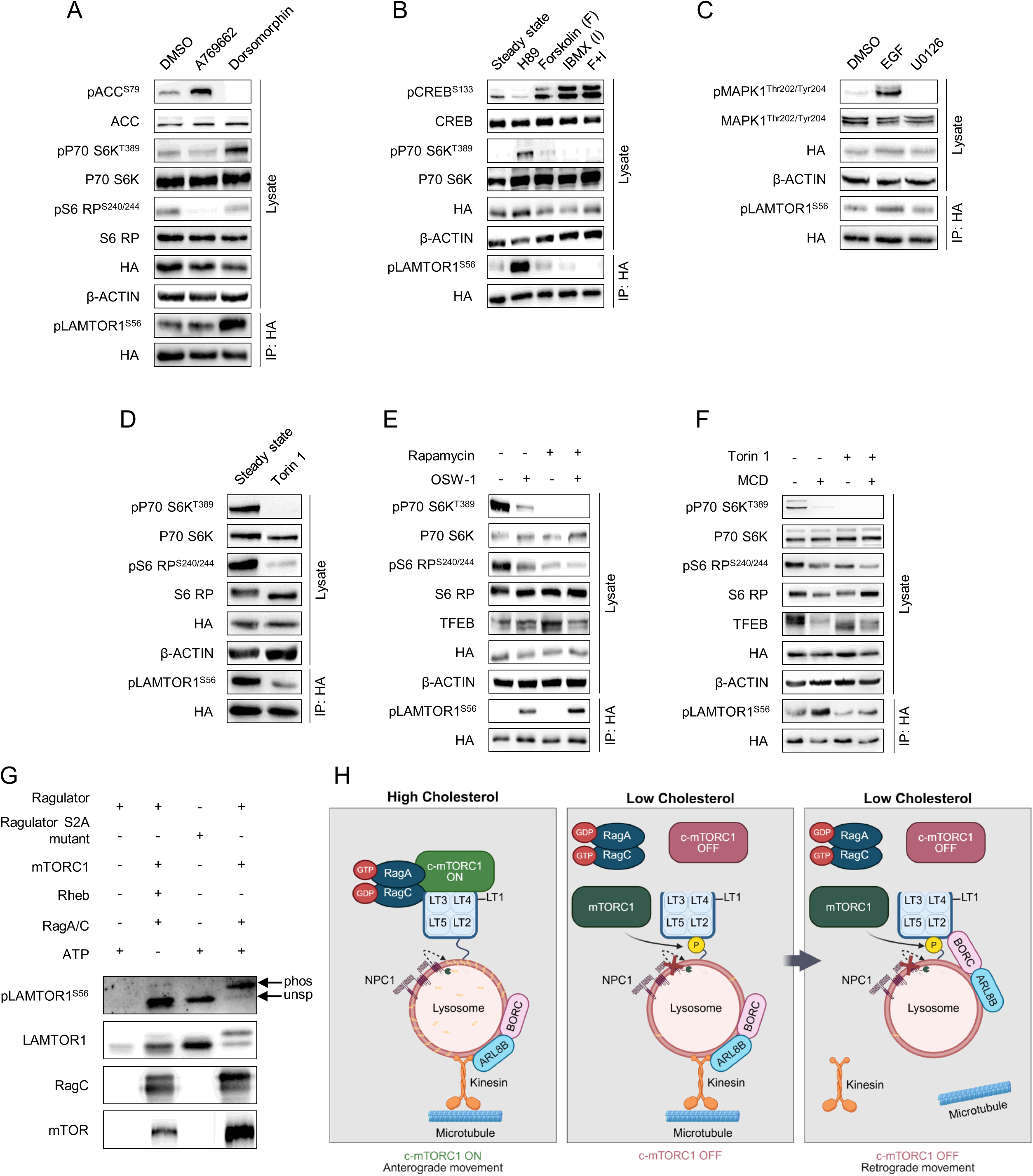
mTORC1 phosphorylates LAMTOR1 pS56. (A) HA-tagged LAMTOR1 WT cells were serum-starved for 2 h and consequently treated with 100 µM A769662 (AMPK activator) or 10 µM Dorsomorphin (AMPK inhibitor) for 1 h. IP eluates and cell lysates were probed with antibodies as indicated. (B) HA-tagged LAMTOR1 WT cells were treated with 15 µM H89 (PKA inhibitor), 10 µm forskolin (F), 500 µM IBMX (I), or F+I (PKA activators) for 24 h. IP eluates and cell lysates were probed with antibodies as indicated. (C) HA-tagged LAMTOR1 WT cells were treated with 40 ng/mL epidermal growth factor (EGF) (MAPK1 activator) and 10 μM of U0126 (MAPK1 inhibitor) for 10 min. IP eluates and cell lysates were probed with antibodies as indicated. (D) HA-tagged LAMTOR1 WT cells were treated with 250 nM Torin 1 (mTOR inhibitor) for 50 min. IP eluates and cell lysates were probed with antibodies as indicated. (E) HA-tagged LAMTOR1 WT cells were treated with 200 nM Rapamycin (mTORC1 inhibitor) for 1 h, 200 nM OSW-1 for 2 h, or a combination of both. IP eluates and cell lysates were probed with antibodies as indicated. (F) HA-tagged LAMTOR1 WT cells were treated with 250 nM Torin 1 for 50 min, 0.5% MCD with 3% LPDS, or a combination of both. IP eluates and cell lysates were probed with antibodies as indicated. (G) mTOR phosphorylates LAMTOR1 at S56. *In vitro* kinase assay using a hyperactive mTOR composite mutant (T1977R + S2215Y on mTOR),^79^ and Ragulator as substrate. Four reactions were performed employing i) Ragulator and ATP; ii) Ragulator, hyperactive mTORC1, RHEB, RagA/C, and ATP; iii) Ragulator S2A mutant (serine residue from the purification tag was mutated to alanine); iv) Ragulator, RagA/C, hyperactive mTORC1, and ATP. Kinase reactions were subjected to immunoblotting against antibodies as indicated. Phos: LAMTOR1 S56 phosphorylation; unsp.: unspecific band. (H) Model for Ragulator complex occupancy regulation by LAMTOR1 S56 phosphorylation. Lysosomal membrane cholesterol depletion induces LAMTOR1 S56 phosphorylation by mTORC1, which displaces the Rag GTPases from the Ragulator complex and facilitates binding of BORC. These events promote deactivation of canonical mTORC1 and retrograde trafficking of lysosomes.

For the regulation of cellular metabolism, AMPK and PKA activities are well known to oppose mTORC1 signaling, reflecting nutrient poor/rich states, respectively. The inverse regulation of LAMTOR1 S56 phosphorylation with all three signaling pathways (AMPK, PKA, and mTORC1) therefore, presents a contradiction. It is by now well established, that mTOR is present in other complexes with distinct substrate specificities, in addition to (canonical) mTORC1, whose activity monitored through P70 S6K and S6 RP phosphorylation was inversely regulated to LAMTOR1 S56 phosphorylation (Figures 2C, 2H, 3C, S2G, S4A, S5I), namely non-canonical mTORC1^12^ and mTORC2.^10^ We, therefore, tested for involvement of mTOR in LAMTOR1 S56 phosphorylation, irrespective of the complex it is present in, by treating cells with Torin 1,^71^ a direct pharmacological inhibitor which binds to the active site of the kinase, and therefore acts on all complexes. Surprisingly, we identified a strong downregulation of LAMTOR1 S56 phosphorylation upon Torin 1 treatment implying a role of mTOR (Figure 5D). As this effect could be due to another downstream effector kinase, we conducted a radioactivity-based *in vitro* kinase screen with a LAMTOR1 peptide harboring S56 for 245 serine/threonine kinases^72^ (Figure S5A, Table S5). Evaluation of the only candidate kinase identified in this assay, mitogen-activated protein kinase kinase kinase kinase 2 (MAP4K2), through NG25-based inhibition^73^ and siRNA-based knockdown, respectively, failed to detect any effect on LAMTOR1 S56 phosphorylation levels (Figure S5B and S5C). Furthermore, with respect to a possible involvement of mTORC2, we identified a downwards-trend for phosphorylation levels of its known substrates in our phosphoproteomics dataset (Figure S5D, Table S4). We confirmed deactivation of mTORC2 upon U18666A treatment by western blot analysis for phosphorylation levels of one of its key substrates, protein kinase B (AKT) S473 (Figure S5E), which is in line with the lacking effect of insulin stimulation (a major activator of mTORC2) on LAMTOR1 S56 phosphorylation observed in our previous experiments (Figure 4B), collectively contradicting the possibility of mTORC2 involvement.

This cumulative evidence motivated us to investigate involvement of a non-canonical version of mTORC1 in LAMTOR1 S56 phosphorylation, as such a complex was reported to directly phosphorylate TFEB with no activity towards the canonical substrates P70 S6K and S6 RP.^12,74,75^ Initially, we ruled out participation of canonical mTORC1 signaling, by combined treatments with OSW-1 and Rapamycin, whose short-term exposure exclusively inhibits canonical mTORC1 signaling through a complex peptidyl-prolyl cis-trans isomerase FKBP1A (FKBP12)^76^ (Figure 5E). In line with our previous findings and the literature, Rapamycin did not show any effect on LAMTOR1 and TFEB,^75^ respectively, further pointing to non-canonical mTORC1 signaling. We then combined MCD and Torin 1 treatment and monitored both LAMTOR1 S56 and TFEB phosphorylation. In agreement with non-canonical mTORC1 signaling, Torin 1 abolished MCD’s effect on LAMTOR1 S56 phosphorylation and, importantly, TFEB phosphorylation dynamics adhered to activation of non-canonical mTORC1 upon MCD treatment (Figure 5F). Additionally, we combined Torin 1 with OSW-1 as well as LB100, a small molecule inhibitor of PP2A which was shown to activate AKT,^77^ further boosting canonical mTORC1 signaling (Figure S5F). In these experiments we observed a Torin 1-related abolishment of LAMTOR1 S56 phosphorylation upon OSW-1 treatment and no effect of PP2A inhibition, further underlining a possible role of non-canonical mTORC1. Taken together, these experiments suggested that LAMTOR1 S56 is phosphorylated by a mTOR version distinct of canonical mTORC1.

Binding of the Rag GTPases to Ragulator is indispensable for canonical mTORC1 activity and non-canonical mTORC1 substrate recognition^12^ and the cryoEM/crystal structures of both complexes^50,51,74^ clearly show that the interaction of RagC and LAMTOR1 occupies the region harboring S56. Furthermore, in all our experiments LAMTOR1 S56 phosphorylation displaced all Rag GTPases from Ragulator (Figure 2C), resulting in disassembly of the mTORC1 nutrient sensing machinery. We, therefore, argued that a Rag GTPase-independent version of mTOR, which is, based on our current evidence distinct from mTORC2, is likely to phosphorylate LAMTOR1 at S56. To further follow up on this, we mis-localized LAMTOR1 to the plasma membrane to uncouple Ragulator from the lysosomal membrane-resident nutrient sensing machinery. Correct lysosomal localization of LAMTOR1 requires N-terminal myristoylation and palmitoylation in combination with a lysosomal sorting motif (LLL) and in yeast, leucine (L) to alanine (A) mutation of this motif was shown to result in mis-localization of LAMTOR1 to the plasma membrane.^78^ Analogously, we generated HA-LAMTOR1 versions with L to A substitutions of L21, L22, and L23 (LLL-AAA). We expressed them in a LAMTOR1 WT background to enable normal lysosomal mTORC1 signaling and to prevent full relocation of the cellular Ragulator pool. We confirmed plasma membrane localization of HA-LAMTOR1-AAA in these cells (Figure S5G) and performed co-IP/-immunostaining experiments for LAMTOR1 with RagB, and mTOR. We identified binding, albeit at low levels, of RagB to plasma membrane-localized HA-LAMTOR1-AAA and co-localization with mTOR (Figure S5H and S5I). Importantly, U18666A treatment of these cells resulted in strong phosphorylation of HA-LAMTOR1-3A at S56 demonstrating independence of the lysosomal membrane-localized nutrient sensing machinery (Figure S5I). Finally, we probed an *in vitro* kinase assay of a recently described hyperactive version of mTORC1^79^ with our LAMTOR1 pS56-specific antibody. Importantly, we observed *in vitro* phosphorylation of LAMTOR1 S56 by mTOR in these experiments (Figure 5G).

Taken together, our findings indicate that mTOR phosphorylates LAMTOR1 at S56 in a rapamycin-insensitive manner as response to reduced cholesterol levels of the lysosomal limiting membrane. LAMTOR1 S56 phosphorylation displaces the Rag GTPases and SLC38A9 from the Ragulator complex, irrespective of amino acid levels or mTORC1 activation by upstream signaling pathways, enabling a cholesterol-specific override of Rag GTPase-mediated inputs. This enables Ragulator to change from a Rag GTPase- to a BORC-bound form, presenting a binding switch for the complex. Binding to Ragulator results in inhibition of BORC, triggering detachment of KIF5B and ARL8B, and hence blockage of BORC-mediated anterograde transport of lysosomes along microtubule tracks.^20,21^ This, consequentially, results in perinuclear immotile lysosomes,^16^ facilitating their engagement in ER-contact sites^8^ for transfer of ER cholesterol to the lysosomal membrane (Figure 5H).

## Discussion

Here, we identify that LAMTOR1 S56 phosphorylation functions as a molecular switch enabling dual use of the Ragulator complex for both mTORC1 signaling and lysosomal positioning. We show that LAMTOR1 S56 phosphorylation is induced exclusively in response to reduced cholesterol levels of the lysosomal limiting membrane and catalyzed by mTOR. The phosphorylation site facilitates displacement of the Rag GTPases from the Ragulator complex and, hence, enables binding of BORC. This results in deactivation of both canonical mTORC1 signaling and BORC-mediated lysosomal anterograde transport. Importantly, LAMTOR1 S56 phosphorylation alone is necessary and sufficient to perform this switch under conditions of amino acid and glucose sufficiency, enabling a cholesterol-specific override of other signals. Consequentially, the mutagenesis-induced prevention of LAMTOR1 S56 phosphorylation resulted in increased binding of Rag GTPases to Ragulator and hyperactive canonical mTORC1 signaling insensitive to perturbations of cholesterol homeostasis and reduced Ragulator-BORC interaction leading to peripheral accumulation of lysosomes. This establishes LAMTOR1 S56 phosphorylation as a regulatory switch for mTORC1 deactivation and lysosomal retrograde movement under reduced cholesterol levels and provides a mechanistic explanation for the dual use of the Ragulator complex by both mTORC1 and BORC.

For our multi-OMICs experiments, we utilized U18666A to inhibit NPC1. It was shown previously, that U18666A can also bind to other proteins, such as oxidosqualene cyclase and desmosterol reductase,^26^ implying the possibility of off-target effects in our MS data sets. While we showed through orthogonal experiments that LAMTOR1 phosphorylation indeed responds only to lysosomal membrane cholesterol levels, we cannot exclude that other regulatory patterns of our datasets, such as upregulation of proteins related to cholesterol biosynthesis, are due to off-target effects of U18666A. Therefore, additional experiments are needed to unequivocally assign individual events to altered lysosomal cholesterol homeostasis.

With respect to differential membrane protein phosphorylation, we identified several other phosphosites on lysosomal proteins to be co-regulated with LAMTOR1 S56, such as VAMP8 T54. This site was shown previously to be involved in lysosome-autophagosome fusion,^42^ a process which is also impaired upon lysosomal cholesterol accumulation, indicating other regulatory mechanisms through phosphorylation of lysosomal surface proteins in response to altered cholesterol homeostasis. Additionally, we identified differentially regulated phosphorylation sites on autophagosomal proteins opening up the possibility of a bidirectional regulatory mechanism involving both organelles, and on the ER and mitochondria. Given the role of lysosome-ER/-mitochondria contact sites for the exchange of cholesterol and other lipids, the fact that we identified differentially phosphorylated residues also on proteins located on the cytosolic face of both organelles, and that we detected differentially regulated lipid species which are directly related to them (acylcarnitines and cholesteryl esters), implies the existence of additional phosphorylation-based regulatory mechanisms in response to altered lysosomal cholesterol homeostasis.

The LAMTOR1 S56 phosphorylation-specific displacement of mTORC1 components from Ragulator enables binding of BORC, whose role for lysosomal positioning is well-established and whose regulation in the context of amino acid level perturbations were investigated in detail.^20,21^ Importantly, overexpression of BORC was demonstrated to displace the Rag GTPases from Ragulator,^53^ underlining the mutually exclusive binding of Ragulator to either BORC or the Rag GTPases, which we also observed in our experiments. The co-existence of LAMTOR1 S56 phosphorylation-mediated Ragulator-BORC complexes with non-canonical mTORC1 signaling occupied Ragulator can be explained by the low stoichiometry of BORC, which is markedly lower (approx. 6-fold) than Ragulator/Rag GTPase levels.^20,21^ This low abundance also necessitated for our studies the use for overexpression systems for investigating the relationship of LAMTOR1 and BORC, as the currently available antibodies for BORC subunits are not sensitive enough, presenting a limitation of this study. We were only able to detect differentially regulated endogenous levels of BORC in our (phospho)-proteomics data of LEFs, which contain several downregulated phosphorylation sites on BORC components. As BORC remains attached to the lysosomal surface also in its inactive form, it is not possible to draw any conclusions based on protein level data obtained from U18666A LEFs. However, we did identify reduced levels of ARL8B, which is crucial for BORC-mediated lysosomal positioning, and who are indicative of its binding to Ragulator. Further, knock down of either ARL8B or BORCS5 was sufficient to rescue the lysosomal distribution phenotype observed for the LAMTOR1 S56A version. For a detailed characterization of Ragulator-BORC binding in response to LAMTOR1 S56 phosphorylation, however, further detailed analyses will be necessary.

It is crucial to mention that binding of Ragulator to BORC results in BORC inactivation, explaining the lack of lysosomal anterograde movement, but providing no explanation for the mechanisms regulating lysosomal retrograde trafficking, and hence perinuclear localization, which we and others observed for altered cholesterol homeostasis.^22^ While several complexes were shown to date to regulate lysosomal trafficking,^80^ a possible player involved in this context could be TMEM55B, as it was demonstrated to mediate retrograde lysosomal transport via JIP4 in response to amino acid starvation and cholesterol-induced lysosomal stress through treatment with U18666A.^22^ In how far LAMTOR1 S56 phosphorylation affects this process, or if a co-regulated phosphorylation of another lysosomal membrane protein is involved, is currently unclear.

A striking feature of LAMTOR1 S56 phosphorylation is the displacement of Rag GTPases, and hence the inactivation of canonical mTORC1, also under levels of nutrient sufficiency, as the latter typically results in association of their active forms (RagA/B^GTP^, RagC/D^GDP^) to Ragulator.^81^ In order to enable spatial access to phosphorylate LAMTOR1 S56, it is crucial that the RagC/D heterodimer dissociates from Ragulator, as the RagC binding interface blocks this part of LAMTOR1.^50^ That it is possible to phosphorylate LAMTOR1 S56 on Ragulator complexes interacting with Rag GTPases is in agreement with a model presented in two recent studies, where the they constantly transition between the lysosomal surface and the cytosol under conditions of nutrient sufficiency. In this model, nutrient levels affect the Rag GTPases residence time on Ragulator, and hence determine the strength of mTORC1 signaling,^15,82^ providing free Ragulator complexes between cycles that enable LAMTOR1 S56 phosphorylation. Importantly, all nutrient sensing mechanisms described so-far impinge on the Rag GTPases, implying that displacement of Rag GTPases by LAMTOR1 pS56, could override all other cellular nutrient inputs for mTORC1.

It was shown recently that also lysosomal membrane cholesterol levels are communicated to mTORC1 via the Rag GTPases. Lysosomal cholesterol is sensed by LYCHOS, which sequesters GTPase-activating protein (GAP) activity toward Rags 1 (GATOR1) in an inactive form at the lysosomal surface under conditions of cholesterol sufficiency. Upon low membrane cholesterol levels, LYCHOS releases GATOR1, which binds KICSTOR in its active form at the lysosomal surface, exerting its GAP function on RagA/B, resulting in their GDP loaded, and hence inactive form.^13^ This connects lysosomal membrane cholesterol levels with modulation of Rag GTPase residence time, providing a possible explanation for the regulation of LAMTOR1 accessibility, and hence S56 phosphorylation, possibly in a negative feedback loop downregulating canonical mTORC1 activity under low lysosomal membrane cholesterol levels.

Importantly, it was shown that RagA/B are crucial for lysosomal recruitment of mTORC1,^12^ assigning their deactivation in this context a role as initiating step, followed by phosphorylation of LAMTOR1 at S56 which prevents re-attachment of RagC/D that in turn results in displacement of all Rag GTPases, and deactivation of mTORC1. With respect to Folliculin-Folliculin-interacting protein (FLCN-FNIP),^23^ which presents the GAP for the RagC/D heterodimer and directly transmits amino acid level information to mTORC1 independent of other nutrient sensing components,^12^ to our knowledge, no connection to cholesterol was demonstrated so far. Cholesterol insensitivity of this pathway could be a possible explanation for our observation of active non-canonical mTORC1^75^ signaling under conditions resulting in LAMTOR1 S56 phosphorylation (based on TFEB phosphorylation patterns^75^ and lack of CLEAR network expression^35^ induction in our proteomics dataset). A possible explanation for the insensitivity of non-canonical mTORC1 could lie in its dependence of FLCN-FNIP, as it promotes GDP-bound RagC/D, which strongly interacts with Ragulator, and therefore sterically blocks phosphorylation of LAMTOR1 S56. Considering that our data strongly suggest that LAMTOR1 S56 is phosphorylated by mTOR, this would imply that this phosphorylation site is at the core of a mTORC1-specific negative feedback loop with no (or only minor) effects towards non-canonical mTORC1.

The fact that canonical mTORC1, through Rag GTPase-mediated recruitment of RAPTOR, and non-canonical mTORC1, through direct interaction of the Rag GTPases with TFEB, both depend on the Rag GTPases for substrate recognition implies that a version of mTORC1 phosphorylates LAMTOR1 which is not bound by the currently known substrate recognition mechanisms. Based on our data, we can exclude a contribution of mTORC2, and a mTORC1-centric negative feedback loop is a likely scenario. We cannot, however, exclude the presence of an unrelated version of mTOR and detailed investigation will be necessary to address this highly interesting question in future studies.

## Acknowledgments

We would like to thank Pineda Antibody Service (Berlin, Germany) for generating the anti LAMTOR1 pS56 antibody, which made this study possible. We are grateful to Madhanagopal Anandapadamanaban and Roger L.William (MRC-LMB, Cambridge, United Kingdom) for providing the invitro kinase assay samples. We also thank Renk Asik and Katharina Baum (Institute for Informatics, Free University Berlin) for performing the GSEA analysis of the proteomic dataset. P.M. received funding from the Jürgen Manchot scholarship, A.D., F.A. and S.F.J. from the German Academic Exchange Service (DAAD), and D.W. received funding from the German Research Council (DFG) as part of FOR2625.

## Author contributions

P.M. performed cell culture, optimization of treatment conditions, and other molecular and cell biology experiments, with the contribution of A.D., A.M., M.B., and S.F.C. A.M. generated HEK NPC1 KO cells. F.A. and N.R. performed TLC analyses. A.D. conducted proteomic and phosphoproteomic analyses of WCL, and co-IPs. P.M. performed lysosome enrichment, and its proteomic and phosphoproteomic analyses. C.C. and R.A. performed lipidomics experiments. M.E. and V.H. carried out immunofluorescence experiments in U18666A-treated MEF, HeLa, and U2OS cells including lysosomal positioning analyses. M.E.G. and L.A. generated LAMTOR1 cell lines. S.R. and M.D. provided the LAMTOR1 LLL-AAA mutant and performed immunofluorescence experiments on these cells. P.M. and D.W. conceptualized the study with input from V.G.; P.M. wrote the original draft; D.W. rewrote and edited the manuscript. All authors reviewed the manuscript and contributed to the final version of the manuscript.

## STAR METHODS

Detailed methods are provided in the online version of this paper and include the following:

- KEY RESOURCE TABLE
- RESOURCE AVAILABILITY

- Lead contact
- Materials availability
- Data and code availability
- EXPERIMENTAL MODEL AND SUBJECT DETAILS

- Cell lines
- METHOD DETAILS

- SILAC and other treatments
- Cell lysis
- Enrichment of lysosomes using SPIONs
- Mass spectrometry sample preparation
- Phosphopeptide enrichment
- LC-MS/MS
- Data processing
- Lipid and metabolite extraction
- Lipid analysis, direct infusion MS and MS/MS analysis
- Metabolite analysis and LC-MS/MS
- Cloning and site-directed mutagenesis
- Transfection and generation of stable cell lines
- CRISPR/Cas9-mediated NPC1 KO cell line generation using HEK293 cells
- Sodium dodecyl sulfate-polyacrylamide gel electrophoresis and western blotting
- Immunoprecipitation
- Immunofluorescence and microscopy
- Acetyl-Coenzyme A assay
- Radioactive protein kinase screening assay
- *In vitro* kinase assay
- Cell perturbations

- Preparation of LPDS
- LPDS treatment
- MCD treatment
- Amino acid starvation
- U18666A treatment
- OSW-1 treatment
- Thin layer chromatography
- Quantification and statistical analysis of western blots

### KEY RESOURCE TABLE

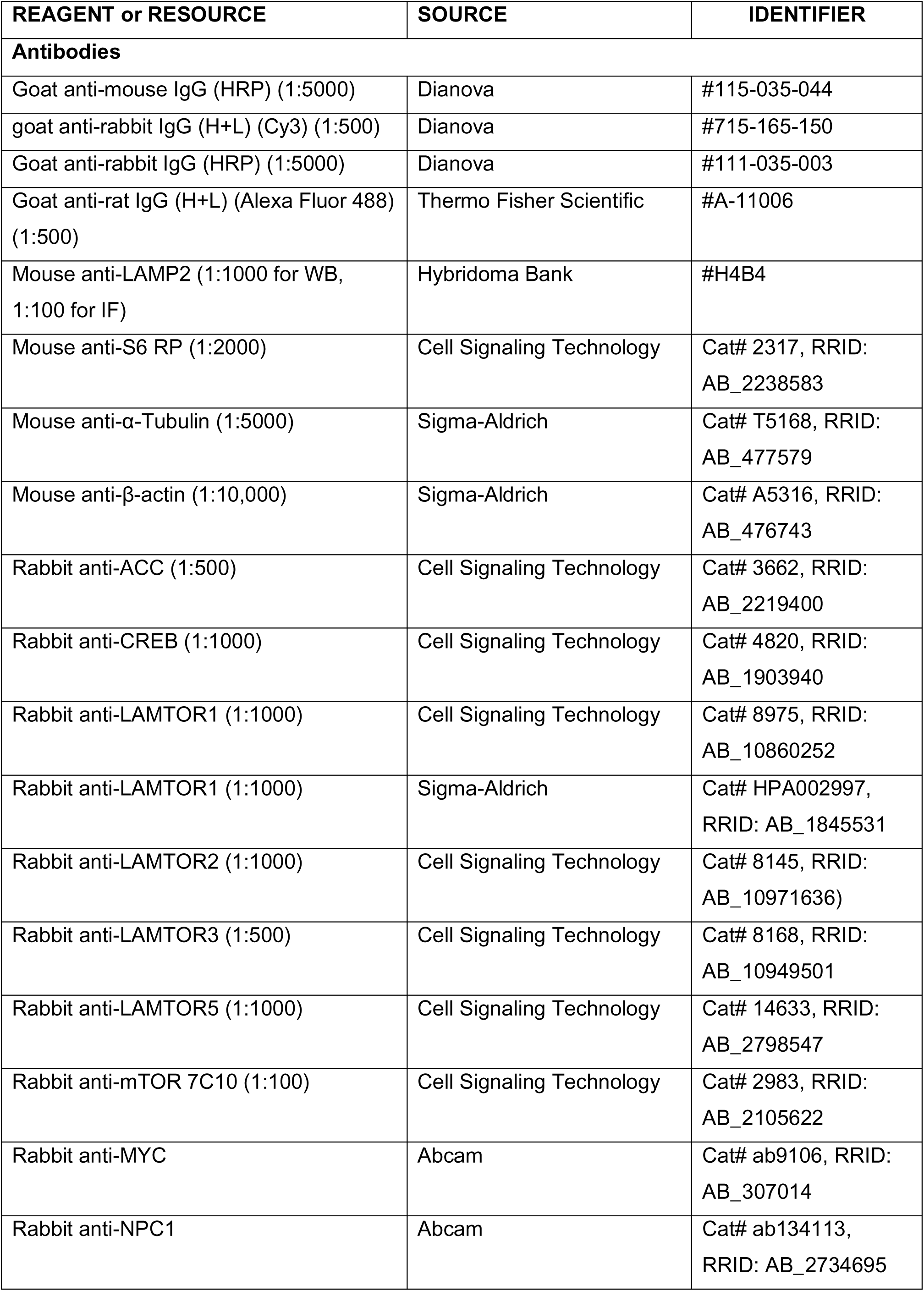
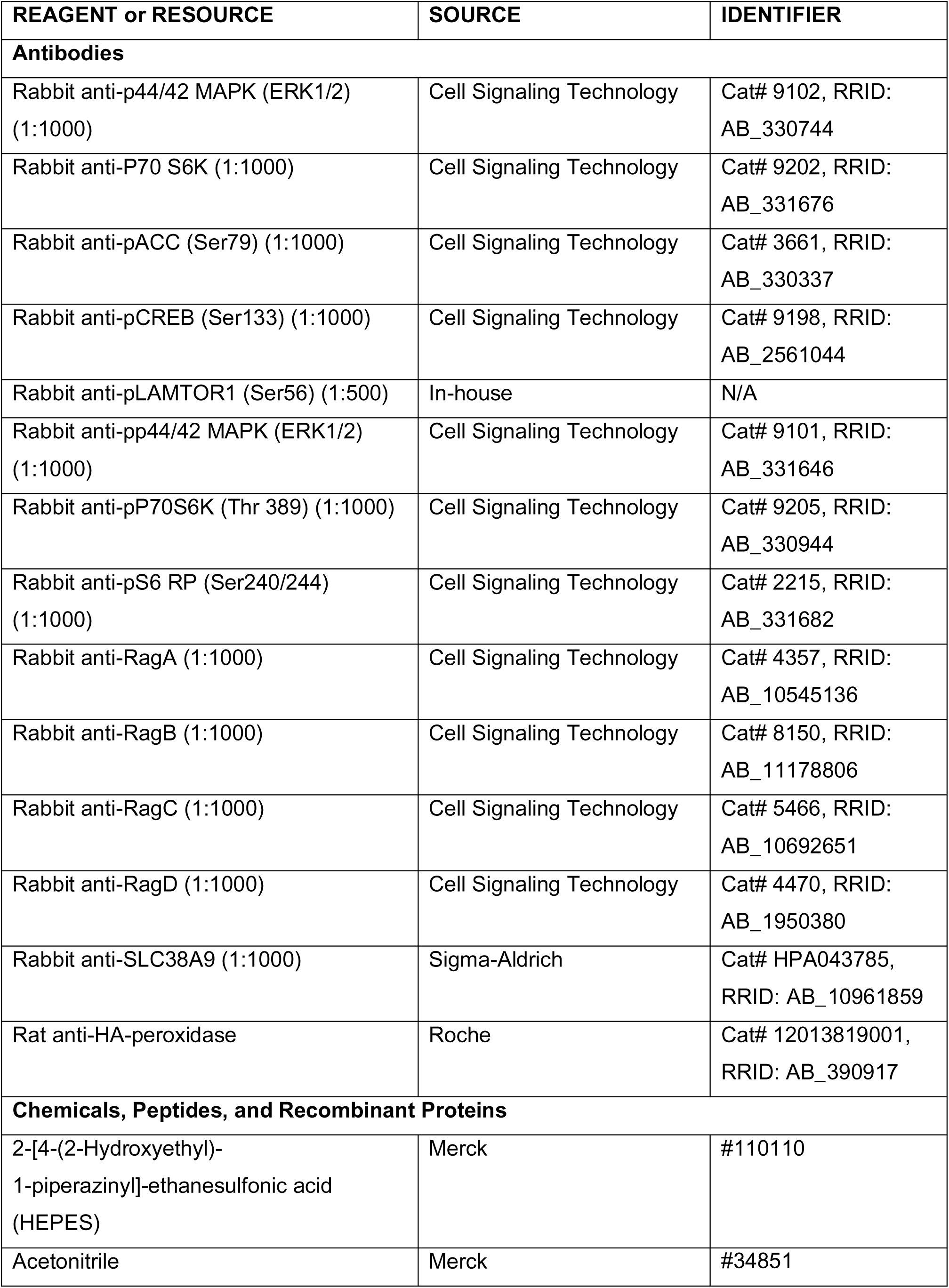
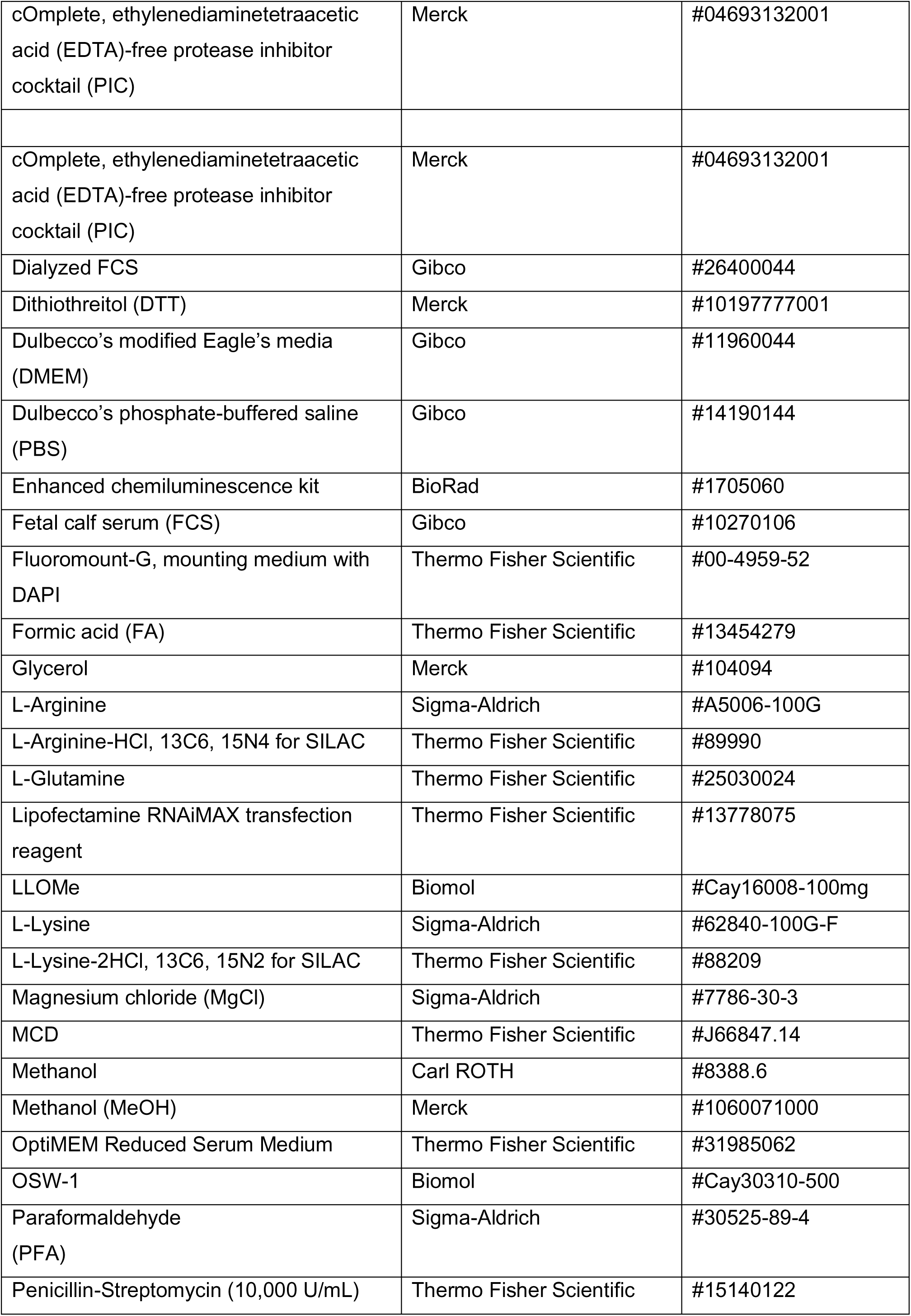
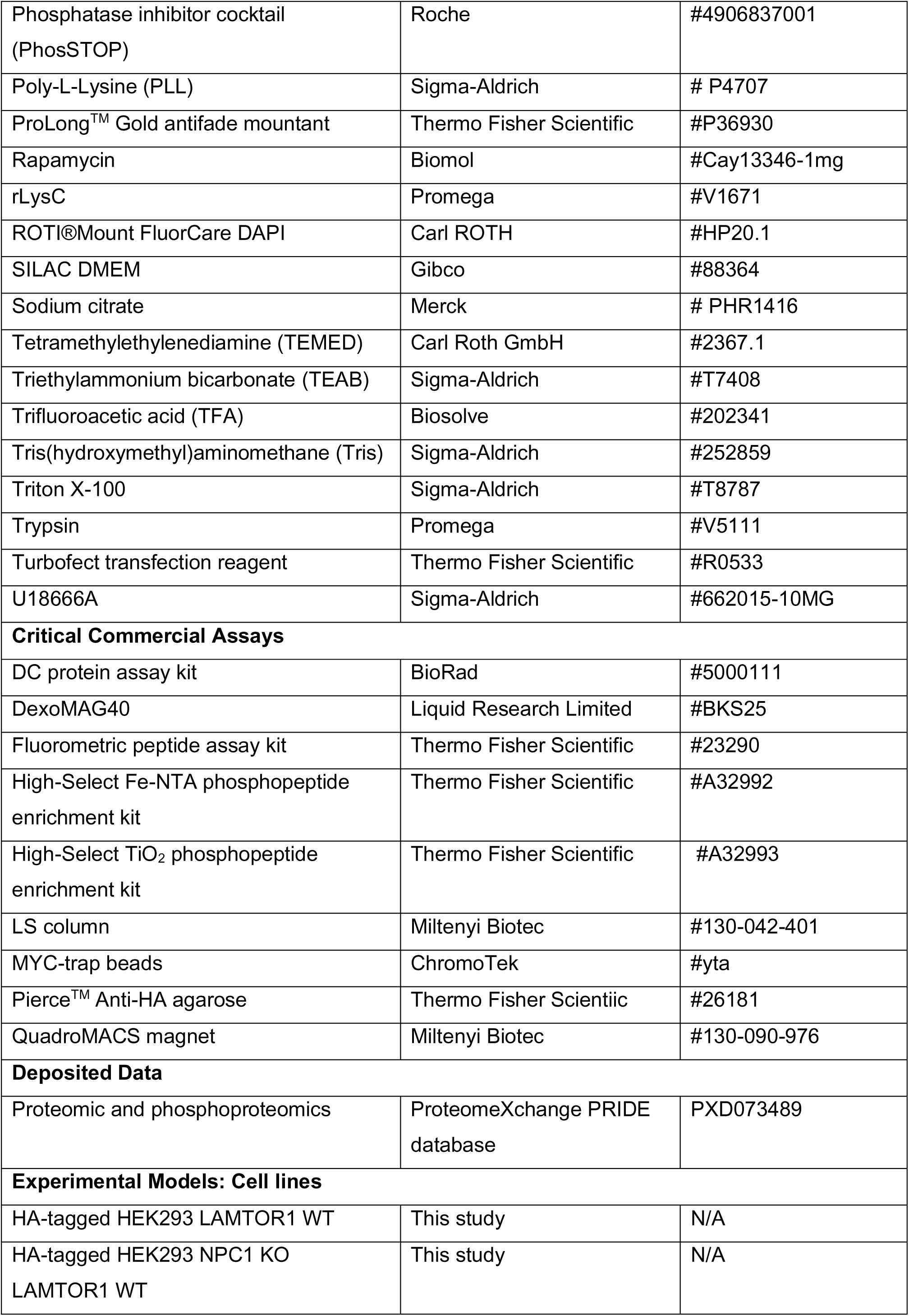
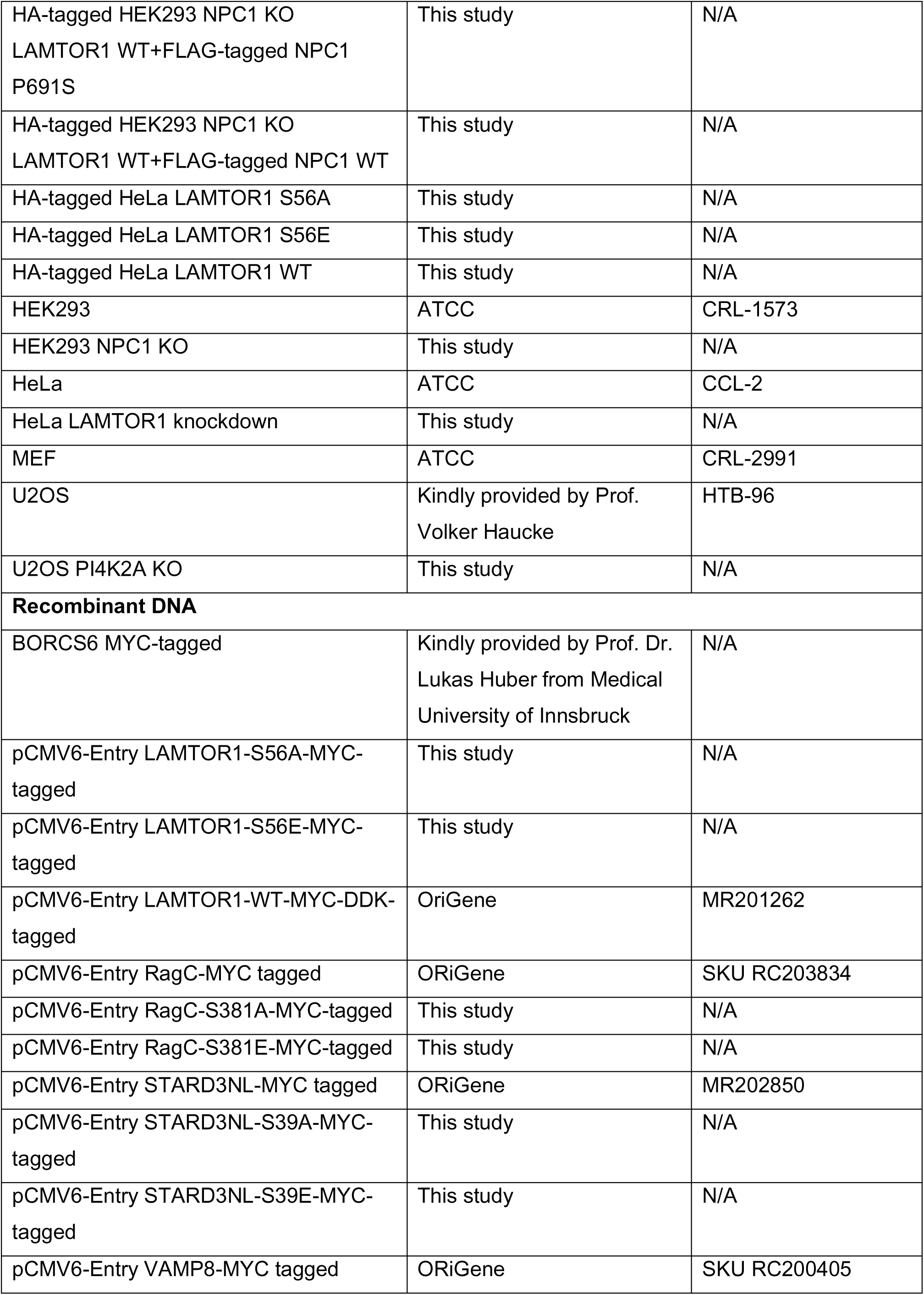
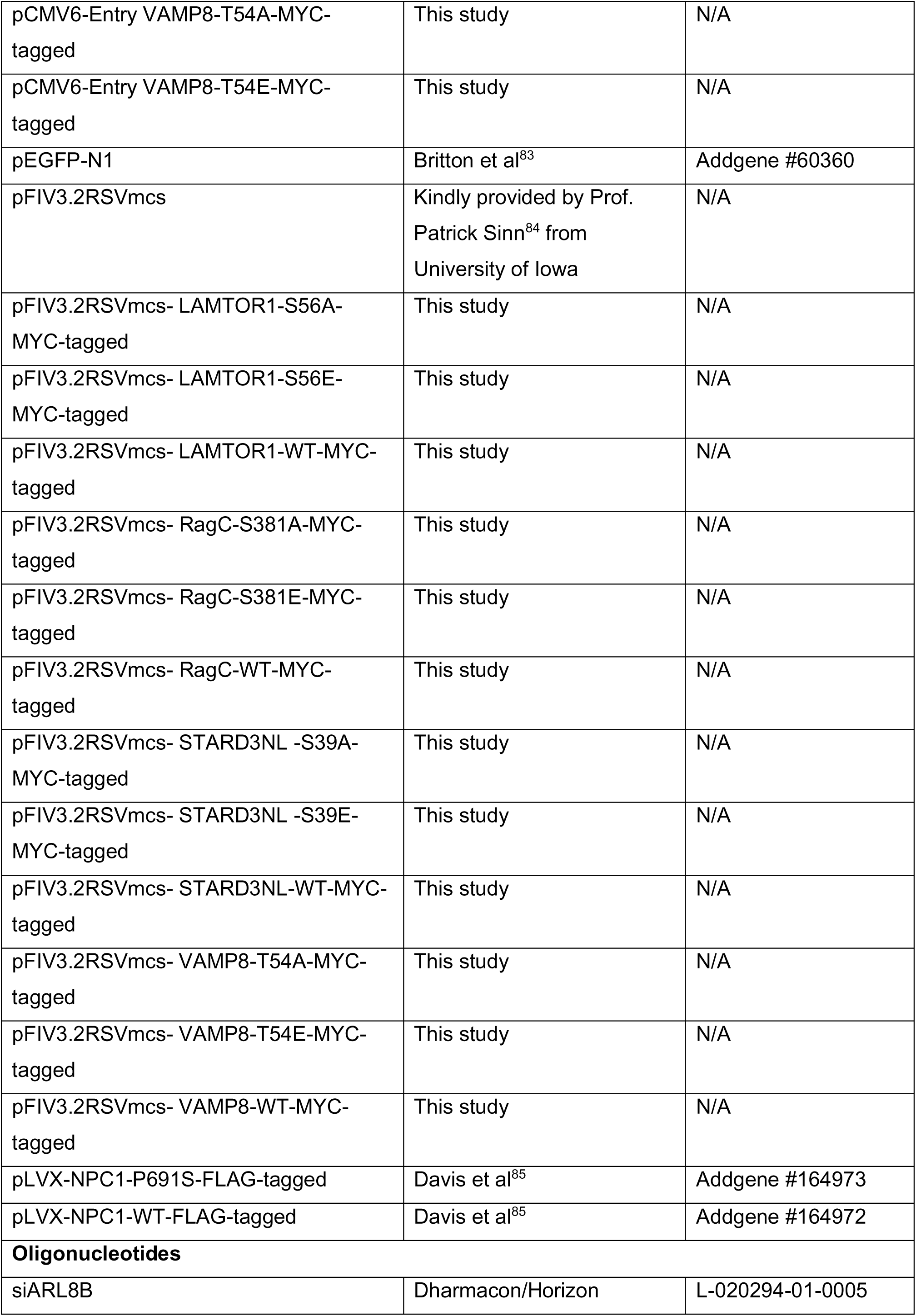
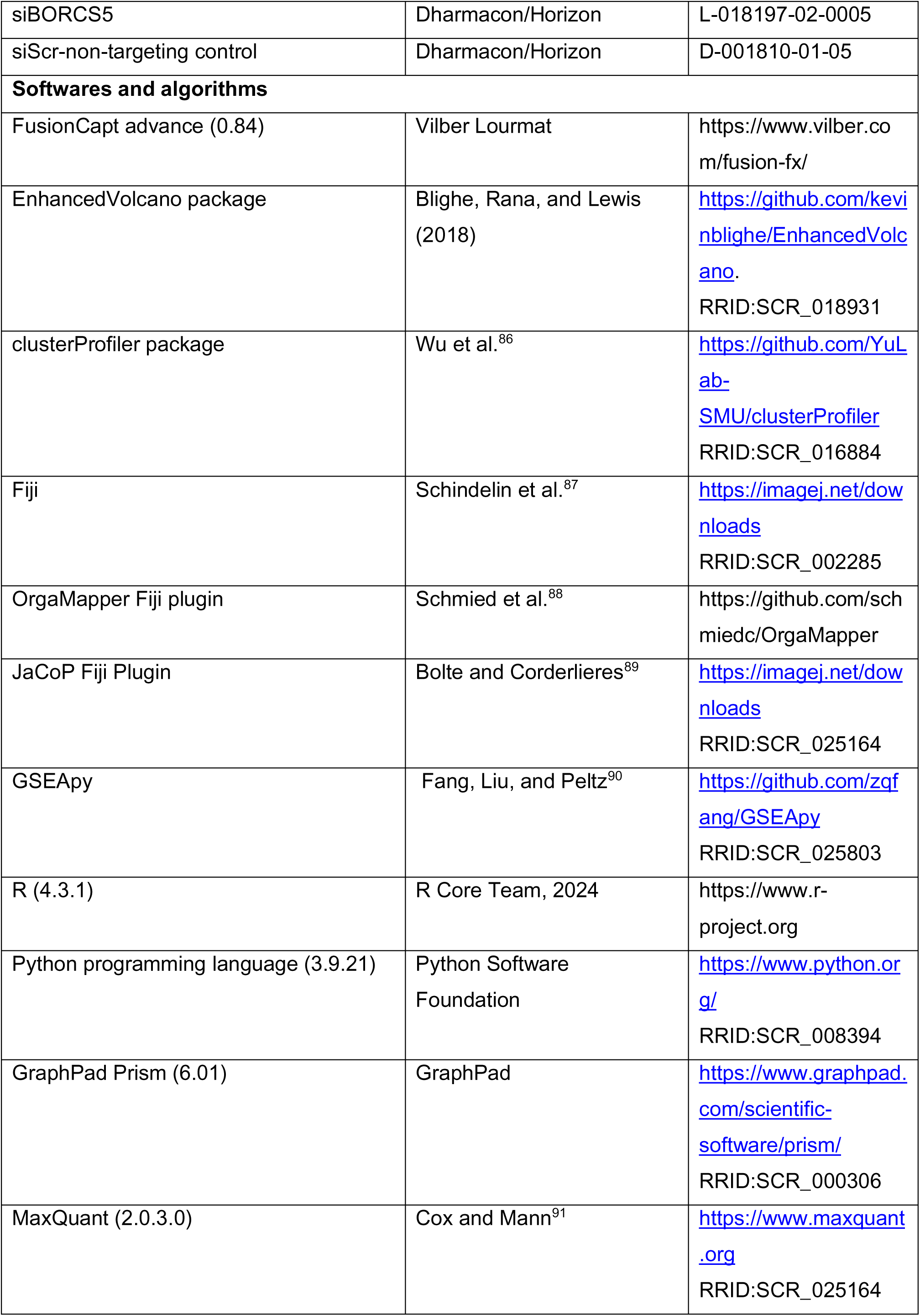
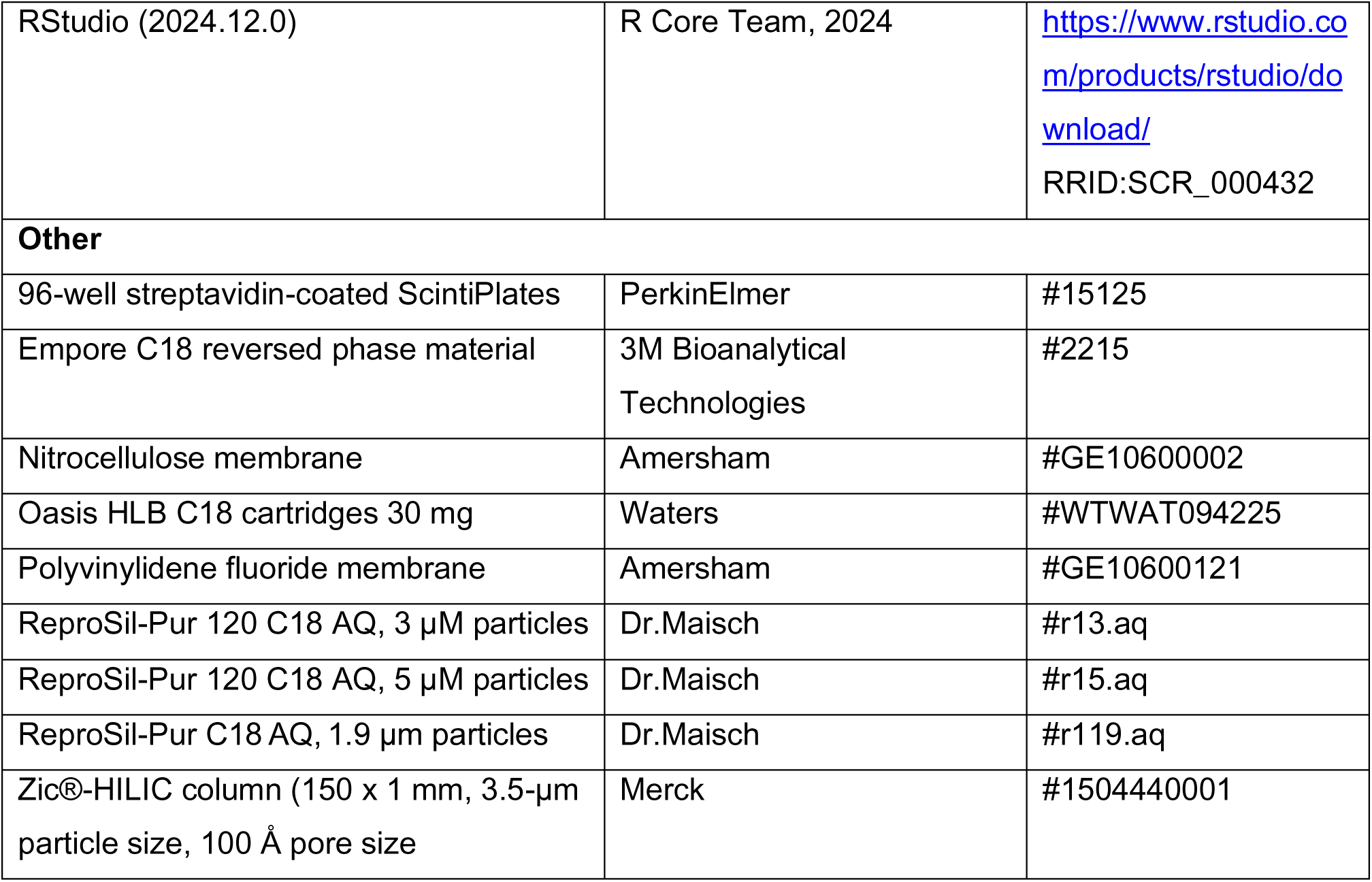

### RESOURCE AVAILABILITY

#### Lead contact

- Further information for resources and requests should be directed to and will be fulfilled by the lead contact, Dominic Winter.

#### Materials availability

- All the unique reagents generated in this study are available from the lead contact without restriction.

#### Data and code availability

- The mass spectrometry proteomics and phosphoproteomics data have been deposited to the ProteomeXchange Consortium via the PRIDE^92^ partner repository with the dataset identifier PXD073489. The lead contact will share all the data reported in this paper upon request.

### EXPERIMENTAL MODEL AND SUBJECT DETAILS

#### Cell lines

HeLa, HEK293, MEF and U2OS cells were obtained from ATCC. They were cultured in DMEM supplemented with 10% heat-inactivated FCS, 2 mM L-glutamine, 100 µg/mL penicillin and 100 µg/mL streptomycin at 37 °C in a humidified incubator with 5% CO_2_. All cell lines were regularly tested and confirmed to be mycoplasma-free by polymerase chain reaction (PCR).

### METHOD DETAILS

Refer to the key resources table for antibodies, chemicals, plasmids, recombinant DNA, oligonucleotides, and software.

#### Cell culture, SILAC labeling, and other treatments

For SILAC labeling, MEF cells were cultured in SILAC DMEM lacking lysine and arginine supplemented with 10% dialyzed FCS, 2 mM L-glutamine, 100 µg/mL penicillin, 100 µg/mL streptomycin and either “Heavy” or “Light” amino acids. Heavy amino acids, Lysine (^13^C_6_/^15^N_2_; Lys8) and Arginine (^13^C_6_/^15^N_4_; Arg10) were added at a final concentration of 181.2 mg/L and 87.8 mg/L, respectively. Light amino acids, Lysine and Arginine were added at a final concentration of 116.6 mg/L and 69 mg/L, respectively. Cells were cultured in the respective amino acids for 5-6 doublings to ensure efficient incorporation of the labeled amino acids.

The following treatments were performed using the indicated concentration, U18666A (3 µg/mL), OSW-1 (200 nM), 0.5% MCD combined with 3% of LPDS, Torin 1 (250 nM), Rapamycin (200 nM), A769662 (100 µM), Dorsomorphin (10 µM), EGF (40 ng/mL), U0126 (10 µM), Forskolin (10 µM), 3-Isobutyl-1-Methylxanthine (500 µM), and H89 (5 µM).

#### Cell lysis

Cells were washed with ice-cold 1x PBS, harvested using a cell scraper, and centrifuged for 2 min at 2000 × g, 4 °C. Unless indicated otherwise, lysis was performed using the following protocol.

Lysis buffer containing 40 mM HEPES pH 7.4, 1% Triton X-100, 2.5 mM MgCl_2_, 10 mM β-glycerol phosphate, 10 mM sodium pyrophosphate, 1 mM sodium fluoride, 1 mM sodium orthovanadate, 2x EDTA-free PIC, and 1x PhosSTOP was added to the cell pellets. Cells were homogenized by passing through a 25-gauge needle 3-5 times, placed on ice for 30 min, and centrifuged for 10 min at 20,000 × g, 4 °C. Following centrifugation, supernatants were transferred to a pre-cooled tube, and protein concentration was determined using the DC protein assay.

A different lysis protocol was adapted to conduct proteomic and phosphoproteomic analysis of WCL from MEF.^93^ Lysis buffer containing 50 mM Tris-HCl pH 8.0, 8 M urea, 75 mM NaCl, 1 mM sodium fluoride, 1 mM sodium orthovanadate, 1 mM β-glycerophosphate, 10 mM sodium pyrophosphate, 1x PIC and 1x PhosSTOP were added to the MEF cell pellets. Cells were homogenized by resuspending in the ice-cold lysis buffer. Homogenates were briefly sonicated (2 pulses of 15 s, 50% amplitude) using a 2 mm probe and centrifuged for 15 min at 5000 × g, 4 °C. Following centrifugation, supernatants was transferred to a fresh tube and protein concentration was determined using the DC protein assay.

#### Enrichment of lysosomes using SPIONs

For this purpose, 10 cm dishes were coated with 0.5 mg/mL PLL dissolved in PBS for 20 min at 37 °C. After coating, dishes were washed three times with PBS. MEF cells were seeded at a density of 3 x 10^6^ cells per PLL-coated dish and were cultured for 24 h in DMEM supplemented with 10% FCS. Lysosome enrichment was performed as described previously.^31^ After 24 h, cells were supplemented with 10% of SPIONs solution (Dextran: 40 kDa, core size: 8 nm, hydrodynamic size: 50 nm, Fe content: 10 mg/mL, solvent: dH_2_O) and were incubated for 24 h at 37 °C. Later, cells were washed with PBS and fresh DMEM containing 10% FCS was added followed by incubation for 24 h at 37 °C. After incubation, cells were washed using ice-cold PBS. Subsequently, cells were harvested in 2 mL isolation buffer containing 250 mM sucrose, 10 mM HEPES pH 7.4, 1 mM CaCl_2_, 1 mM MgCl_2_, 1.5 mM MgAc, 10 mM sodium pyrophosphate, 10 mM β-glycerophosphate, 1 mM sodium orthovanadate, 50 mM sodium fluoride, 2x cOmplete EDTA-free PIC, and 1x PhosSTOP.

A total of eight dishes of cells were used per condition. Cells belonging to the same condition from eight dishes were pooled into a 15 mL Douncer homogenizer tube and were dounced using 30 strokes. Subsequently, cell homogenate was transferred to a fresh 15 mL tube and was centrifuged for 10 min at 500 × g, 4 °C. After centrifugation, post-nuclear supernatant (PNS) was collected, transferred to a new tube, and the remaining cell pellet was resuspended again in 4 mL of isolation buffer. This cell homogenate was dounced again using 30 strokes, centrifuged, and the resulting PNS was combined with the previously obtained fraction. This fraction was used as the input for lysosome enrichment. Two LS columns were used per lysosome enrichment along with a QuadroMACS magnet. At first, LS columns were equilibrated using 0.5% BSA in PBS at room temperature (RT). The PNS fraction was divided into two halves and each half was passed through one LS column, flow through was collected and LS columns were washed three times using isolation buffer. Further, 1 mL of isolation buffer was added to each LS column and a plunger was employed to elute lysosomes. Lysosomal yield and purity were determined using β-hexosaminidase and DC Protein Assays. The eluted lysosomes from two LS columns belonging to the same condition were pooled and centrifuged for 30 min at 20,000 × g, 4 °C. The resulting lysosomal pellet was further utilized for mass spectrometric analyses.

#### Mass spectrometry sample preparation

The protocol for in-gel digestion was adapted from Shevchenko et al.^94^ 100 µg of MEF WCL along with SDS sample buffer (without reducing agent) was heated at 95 °C for 10 min, resolved on a 10% SDS-PAGE gel, and stained using Coomassie Blue for 1 h at RT. Subsequently, the gel was washed using ddH_2_O, and each lane was cut into small pieces, which were further cut into 1 mm cubes using a scalpel. The gel pieces were destained using 30% ACN in 0.07 M ammonium bicarbonate (NH_4_HCO_3_). Protein disulfide bonds were reduced for 30 min at 600 rpm, 56 °C using DTT (5 mM final concentration). Free SH groups were alkylated using acrylamide (20 mM final concentration) for 30 min at 600 rpm, RT.^95^ The alkylation reaction was quenched by adding DTT (5 mM final concentration), and incubating for 15 min at RT. The gel pieces were washed using 0.1 M NH_4_HCO_3_, dehydrated by adding 100% ACN, dried in a vacuum centrifuge, and incubated in trypsin overnight at 37 °C. After tryptic digestion, the supernatant containing peptides was carefully transferred to fresh tubes. Residual peptides from the gel were collected by consecutively adding 0.1% TFA/50% ACN, 0.1 M NH_4_HCO_3_, and 100% ACN. All the peptide extracts were dried using a vacuum centrifuge and desalted using C18 StageTips.^96^ The desalted peptides were again dried using a vacuum centrifuge and resuspended in 5% ACN and 5% FA.

For in-solution digestion of MEF WCL, protein disulfide bonds were reduced using DTT (5 mM final concentration) for 30 min at 600 rpm, 56 °C, and SH groups were alkylated using acrylamide (20 mM final concentration) for 30 min at 600 rpm, RT.^95^ The alkylation reaction was quenched by adding DTT (5 mM final concentration), and incubating for 15 min at RT. Next, the solution was diluted to 1:5 using 25 mM Tris-HCl (pH 8.2) followed by addition of CaCl_2_ at a final concentration of 1 mM. Subsequently, trypsin was added at an enzyme-protein ratio of 1:200 and the samples were incubated overnight at 37 °C. The following day, peptides were collected, dried in a vacuum centrifuge, and desalted using 10 mg Oasis HLB C18 cartridges. The desalted peptides were dried once more using a vacuum centrifuge.

For in-solution digestion of MEF LEF, the pellets from the SPIONs-enriched lysosomal fractions were resuspended in 8 M urea in 0.1 M TEAB. Equal protein amounts of light and heavy samples were taken; disulphide bonds were reduced, SH groups were alkylated, and reactions were quenched as described previously.^95^ Next, samples were diluted 1:1 using 0.1 M TEAB and incubated with rLysC at an enzyme:protein ratio of 1:200 for 6 h at 1,200 rpm, 37 °C. Later, the final concentration of urea was adjusted to 1.3 M, trypsin was added at an enzyme:protein ratio of 1:100, and the samples were incubated overnight at 1,200 rpm, 37 °C. The following day, peptides were collected, dried using a vacuum centrifuge, and desalted using 10 mg Oasis HLB C18 cartridges.

#### Phosphopeptide enrichment

Phosphopeptide enrichment using MEF WCL was performed as previously reported.^93^ The peptides were resuspended in 80% ACN, 5% TFA and 1 M glycolic acid. TiO_2_ bead slurry were added to the peptides in a ratio of 1:6 (peptide: TiO_2_ beads) incubated for 15 min at 1,200 rpm, RT. Next, beads were centrifuged for 1 min at 13,000 × g, beads were transferred to a fresh tube containing 80% ACN, 1% TFA and centrifuged again. Thereafter, beads were transferred to 20% ACN, 0.1 TFA solution, centrifuged again to remove the supernatant and beads were dried using a vacuum centrifuge. For elution of phosphopeptides, 1% ammonium hydroxide (NH_4_OH) was added to the beads, centrifuged for 15 min at 1,200 rpm, RT. The peptides were acidified by adding FA to a final concentration of 0.1%, dried in a vacuum centrifuge, and resuspended in 70% ACN/0.1% TFA. Following this, TiO_2_ beads were then added again to the samples at a ratio of 1:6 (peptide: TiO_2_ beads), washed using 50% ACN and 0.1% TFA. Peptides were eluted by adding 1% NH_4_OH to the beads, centrifuged for 15 min at 1,200 rpm, RT. Samples were acidified by adding FA to a final concentration of 0.1%, desalted using 10 mg Oasis HLB C18 cartridges and dried using a vacuum centrifuge.

For phosphopeptide enrichment from MEF LEF, initially High-Select TiO_2_™ phosphopeptide enrichment kit was employed according to the manufacturer’s instructions. The flow through obtained from this enrichment was used as input for a second enrichment using High-Select™ Fe-NTA phosphopeptide enrichment kit as per the manufacturer’s instructions.

For fractionation of phosphopeptides, SCX method was performed. Elution buffers containing 7 mM monopotassium phosphate (KH_2_PO_4_), 30% ACN were prepared using ascending concentrations of potassium chloride (KCl) (0, 30, 60, 90, 120, and 350 mM) and pH was adjusted to 2.65. Tip columns were prepared by placing 12 disks of Empore Cation-SR material into a 200 µL micropipette tip. The tip columns were equilibrated using 100 µL of MeOH, elution buffer (7 mM KH_2_PO_4_, 30% ACN, and 350 mM KCl), water, equilibration buffer (50 mM K_2_HPO_4_, pH 7.5, 500 mM NaCl), water and elution buffer (7 mM KH_2_PO_4_, 30% ACN) in the same order. The columns were centrifuged for 3 min at 2,500 x g, RT. Peptides were solubilized in 7 mM KH_2_PO_4_, 30% ACN and were loaded onto the equilibrated tip columns. The tip columns containing samples were centrifuged at 2,500 x g for 3 min at RT and the first flow through was collected. Consequently, 200 µL of the elution buffers containing ascending KCl concentrations were used for eluting five more fractions by centrifuging for 3 min at 2,500 x g, RT. At the end, all the fractions were dried using a vacuum centrifuge.

#### LC-MS/MS analysis

All the peptide extracts were resuspended using 5% ACN and 5% FA. The peptides and phosphopeptides obtained from WCL were analyzed using a nanoflow UHPLC system (Easy-nLC 1000) coupled to an LTQ Orbitrap Velos mass spectrometer (both Thermo Fisher Scientific). The samples were applied to in-house manufactured columns at a solvent flow rate of 1 µL/min (0.1% FA in H_2_O). The columns were prepared by packing a 360 µm outer diameter/ 100 µm inner diameter fused silica capillaries using a P2000 laser puller (Sutter Instruments) with 5 µm ReproSil-Pur 120 C18-AQ particles. Before measuring phosphopeptides, columns were equilibrated with three washes of 50 mM citrate.^97^ Sample separation was performed at a flow rate of 400 nL/min using a 90 min gradient from 99% solvent A (H_2_O, 5% dimethylsulfoxide (DMSO), 0.1% FA), 1% solvent B (ACN with 5% DMSO, 0.1% FA) to 65% solvent A, 35% solvent B. 240 min segmented gradients were used for the unfractionated samples. The segmented gradients consisted of a 200 min linear gradient from 1 to 25% solvent B, followed by a 30 min step to 30% solvent B and a 10 min step to 35% solvent B at a flow rate of 400 nL/min. At 1.6 kV in positive ion mode, peptides eluting from the column were ionized within the nanosource of the mass spectrometer. Survey scans were obtained within the mass analyzer from 400 to 1200 m/z at a resolution of 30,000 and fragmentation of the top 10 most abundant ions. Singly charged ions and ions with uncharged states were omitted from MS/MS fragmentation. Dynamic exclusion was set to 30 s.

The peptide and phosphopeptide extracts from LEF were analyzed using an Ultimate 3000 RSLCnano systemonline coupled to an Orbitrap Fusion Lumos mass spectrometer (both Thermo Fisher Scientific). The analytical columns were produced in-house as described above with different parameters: 50 cm spray tips were prepared from 360 µm outer diameter/ 100 µm inner diameter fused silica capillaries. The columns were packed with 3 µm ReproSil AQ-C18 particles. The columns were washed with 50 mM citrate before measuring phosphopeptide samples. Samples were loaded onto the column with 100% solvent A (0.1% FA) at a flow rate of 600 nL/min and were separated with linear gradient of 240 min using Data-Dependent Acquisition (DDA) from 5-35% solvent B (95% ACN, 0.1% FA) at a flow rate of 300 nL/min. For DDA analyses, survey scans were acquired in the Orbitrap mass analyzer over a mass range of 375-1,200 m/z, with a resolution of 60,000, a maximum injection time of 50 ms, and an AGC target of 4 x 10^5^. The most abundant precursor ions were fragmented in top-speed mode (cycle time: 3 s) using higher-energy collisional dissociation with a collision energy of 30%. Precursors ions with charge states of +2 to +4 were selected for fragmentation. MS/MS spectra were acquired in the Orbitrap at a resolution of 30,000 with dynamic exclusion of 120 s. The precursor mass isolation window was set to 1.6 m/z, with an AGC target of 5 x 10^4^ and a maximum injection time of 54 ms.

#### Data processing

DDA raw files from MEF WCL and MEF LEF were analyzed using Maxquant 2.0.3 and Maxquant 2.0.6, respectively.^91^ Uniprot mouse (release 2023-07; 17,162 entries) and a common contaminant database^98^ were used for the analyses. MS/MS searches were carried out with the following parameters: Trypsin as the digestion enzyme; precursor ion mass tolerance: 4.5 ppm, Orbitrap fragment ion mass tolerance: 20 ppm, fixed modification: propionamide, variable modifications, oxidation at methionine, acetylation at N-termini, number of missed cleavage sites: 2 and match between runs: ON. Data were exported from Maxquant and filtered for contaminants at 5% FDR on peptide and protein level. Subsequently, data were analyzed using Excel, R studio and visualized using R packages: EnhancedVolcano, and Clusterprofiler. GSEA^99^ was performed on differential protein expression using the GSEApy package.^90^ Proteins were ranked by the product of fold change and absolute log10 p-value in descending order. GSEA was performed separately on GO biological processes and GO cellular components using default settings.

#### Lipid and metabolite extraction

The frozen pellets were placed on ice and the SIMPLEX workflow was employed as previously described.^100^ Briefly, cold MeOH was added to each sample, vortexed for 20 s, and incubated in liquid nitrogen for 1 min. Next, the samples were allowed to thaw at RT and sonicated for 10 min at 4 °C. This procedure was repeated three times. In the next step, cold Methyl-tert-butyl ether was added and the mixture was incubated for 1 h at 4 °C in a thermomixer. Phase separation was induced by adding 188 µL water containing 0.1% ammonium acetate. The extract was centrifuged at 10,000 g for 5 min and the upper phase was collected, dried under nitrogen flow and dissolved in 2-Prop/MeOH/CHCl_3_ (4:2:1, v/v/v) containing 7.5 mM ammonium acetate for shotgun analysis. For protein precipitation, MeOH was added to the remaining lower phase in a final ratio of 4:1, v/v MeOH/H_2_O, and the samples were incubated for 2 h at -80 °C, followed by 30 min centrifugation (13,000 g) at 4 °C. The resulting supernatant was collected and the remaining pellet dissolved in 1% SDS, 150 mM NaCl, 50 mM Tris (pH 7.8). The dried metabolite extracts were resuspended in ACN/H_2_O (9:1, v/v) and then stored at -80 °C prior to LC/MS analysis.

#### Lipid analysis, direct infusion MS and MS/MS analysis

The lipid extracts were placed on a 96-well plate and then infused via the robotic nanoflow ion source TriVersa NanoMate (Advion BioSciences) into a Q Exactive Plus (Thermo Fisher Scientific). The ion source was controlled by Chipsoft 8.3.1 software. Ionization voltage was set to +1.25 kV in positive and -1.25 kV in negative mode, and the back pressure was set at 0.95 psi in both polarities. The S-lens level was 60%, and the ion transfer capillary temperature was adjusted to 250 °C. Polarity switching of the TriVersa NanoMate was triggered by the mass spectrometer via contact closure signal as described previously.^101^

For DDA experiments, full MS spectra was acquired at a target mass resolution Rm/z 200 of 140,000 and the top 20 precursors were selected with a 1 Da window for fragmentation. MS/MS scans were acquired at a target mass resolution R_m/z 200_ of 35,000. All spectra were imported in LipidXplorer into a MasterScan database under the following settings: mass tolerance 5 ppm; m/z range 300-1200 (negative mode) and 350-1200 (positive mode); minimum occupation 0.5; intensity threshold 1 x 10^4^. Lipid identification was carried out as described previously.^102,103^

#### Metabolite analysis and LC-MS/MS

Metabolites were analyzed on an UltiMate 3000 RSLCnano system coupled to a QTRAP 6500 mass spectrometer (AB SCIEX). Separation was carried out on a Zic®-HILIC column (150 x 1 mm, 3.5-µm particle size, 100 Å pore size). The mobile phases were 90% ACN (A) and 20 mM ammonium acetate (pH 7.5) in H_2_O (B). The gradient eluted isocratically with 90% ACN for 2.5 min followed by an increase to 60% over 14 min and held at 60% for 2 min. Subsequent reconstitution of the starting conditions and re-equilibration with 100% A for 10 min resulted in a total analysis time of 35 min. The LC separation was carried out at 25 °C under a flow rate of 100 µL/ min. Electrospray ionization Turbo V source parameters were set as follows: curtain gas, 30 arb. unit; temperature, 350 °C; ion source gas 1, 40 arb. unit; ion source gas 2, 65 arb. unit; collision gas, medium; ion spray voltage, 5500 V/ -4500 V (positive mode/ negative mode). For the selected reaction monitoring mode Q1 and Q3 were set to unit resolution and the transitions for all compounds were adapted.^104,105^ For data acquisition and analysis Analyst (1.6.2) and MultiQuant 3.0 were used respectively.

#### Cloning and site-directed mutagenesis

HA-tag was inserted at the C-terminus of LAMTOR1 WT using Gene sewing. Site-directed mutagenesis was performed to mutate serine 56 to glutamic acid (S56E) and alanine (S56A). LAMTOR1 WT, S56E and S56A genes were PCR-amplified from the donor plasmid and subcloned into the retroviral construct, pQXCIN via recombination. Target cells were co-transfected with the pQXCIN and pVSVG constructs using Lipofectamine LTX. After 48 and 72 h, the viral supernatant was collected and directly used for infecting HeLa LAMTOR1 knockdown cell line. After 48 h, infected cells were collected using 400 µg/mL of geneticin.

#### Transfection and generation of stable cell lines

For Turbofect-mediated transfection, cells were seeded at the density of 75 x 10^4^ per 10 cm dish and were cultured for 24 h prior to transfection. The transfection mix was prepared by mixing 10 µg of DNA with 1 mL of serum-free medium and 20 µL of Turbofect transfection reagent and was incubated for 20 min. After incubation, the transfection mixture was added to the cells dropwise. The cells were supplemented with fresh medium 4 h post-transfection and were incubated for 48 h. Later, the cells were harvested and cell lysates were prepared.

For calcium phosphate-mediated transfection, cells were seeded at the density of 75 x 10^4^ per 10 cm dish and were cultured for 24 h before transfection. For transfection, a calcium phosphate-DNA mixture was prepared by combining 10 µg of DNA with 2 M CaCl_2_.^106^ One volume of this mixture was added to one volume of HEPES-buffered saline while the tubes were tapped to ensure mixing. The calcium phosphate-DNA mixture was immediately added to the cells dropwise. Fresh medium was supplemented to the cells 16 h post-transfection and cells were incubated for another 32 h, after which the cells were harvested and lysed.

#### CRISPR/Cas9-mediated NPC1 KO cell line generation using HEK293 cells

Synthetic CRISPR RNA (crRNA) was designed using Dharmacon™ CRISPR design tool (https://horizondiscovery.com/en/custom-synthesis/custom-synthesis-tools) and a synthetic edit-R trans-activating (tracr)RNA with a Cas9 (PuroR) Expression Plasmid vector from Dharmacon™ were prepared according manufacturer’s instructions. For the generation of NPC1 KO cell line, HEK293 cells were seeded onto a 24-well plate (15 x 10^3^ cells per well) 24 h prior transfection. On the day of transfection, 10 µM tracrRNA, together with 10 µm gene-specific crRNA and Cas9 plasmid (100 ng/µl) were co-transfected using DharmaFECT Duo working solution, together with the respective controls, according manufacturer’s instructions. 48 hours post-transfection cells were selected using puromycin for 2 to 4 days. A limited dilution was performed to singularize cells. Single cell colonies were expanded for 2 to 3 weeks and were divided into three 96 well plates. One 96 well plate with single cell colonies was used for PCR and sequencing. Positive clones were validated using western blotting.

#### Sodium dodecyl sulfate-polyacrylamide gel electrophoresis (SDS-PAGE) and western blotting

Polyacrylamide separating and stacking gels were prepared with 10% SDS (w/v), 40% acrylamide (v/v), 10% ammonium persulfate (w/v) (APS), 1% TEMED. 1.5 M Tris-HCl (pH 8.8) and 0.5 M Tris-HCl (pH 6.8) were used for separating and stacking gels, respectively. The samples were heated at 95 °C for 10 min in 1x SDS-sample buffer (stock composition (4x): 240 mM Tris-HCl (pH 7.4), 4% β-mercaptoethanol (v/v), 8% SDS (w/v), 40% glycerol (v/v), 4% bromophenol blue (w/v)). After heating, the samples were loaded onto the polyacrylamide gels, and gel electrophoresis was performed at 80-120 V for 2 h.

For western blotting, proteins were transferred onto either nitrocellulose or polyvinylidene fluoride membranes using a semi-dry or wet blotter for 1 h or 2 h at 0.2 A per membrane. Next, the membranes were blocked using 5% bovine serum albumin (BSA) (w/v) or 5% skimmed milk (w/v) in Tris-buffered saline containing 0.1% Tween 20 (v/v) (TBST) for 1 h or 2 h at RT followed by incubation with respective primary antibodies overnight at 4 °C. For certain antibodies, the blots were cut and incubated separately in respective primary antibodies. The next day, the membranes were washed with TBST three times, 10 min each and were incubated with individual secondary antibodies for 1 h at RT. Subsequently, the membranes were washed three times with TBST, 10 min each. At last, enhanced chemiluminescence reagent was applied on the membrane, and the protein expression signals were visualized using the FUSION SOLO 4 M system (VilberLourmat), and the signals were analyzed using FusionCapt advanced software.

For stripping of blots, blots were incubated in 25 mM glycine (pH 2.0), 1% SDS (w/v) for 10 min at RT. Blots were washed twice with 1x PBS for 10 min. Subsequently, blots were washed using TBST three times for 5 min each. Following that, the blots were blocked using respective blocking buffer and were incubated with primary antibody.

#### Immunoprecipitation

Cells were rinsed with ice-cold PBS twice and were harvested using a cell scraper in ice-cold PBS. The cells were resuspended in lysis buffer containing 50 mM Tris-HCl (pH 8.0), 120 mM NaCl, 1 mM EDTA, 1% Triton X-100, 10 mM pyrophosphate, 10 mM β-glycerophosphate, 2x PIC and 1x PhosSTOP. Cell lysates were homogenized by passing the cells through 25-gauge needle and were incubated for 30 min on ice. Later, the samples were centrifuged for 10 min at 20,000 × g, 4 °C. The supernatant was collected in a pre-cooled tube and the protein concentration was determined using DC protein assay.

For HA-IP, 1 mg of cell lysate was incubated with 20 µL of HA agarose bead slurry in the dilution buffer (150 mM NaCl, 10 mM Tris-HCl (pH 8.0), 0.5 mM EDTA, 1x PIC) overnight at 4 °C in an end-over-end rotator. After incubation, the beads were centrifuged for 2 min at 2,500 × g, 4 °C. After IP incubation, the beads were centrifuged for 2 min at 2,500 × g, 4 °C and were washed with dilution buffer three times. At the end, proteins were eluted from the beads by heating the beads in 2x SDS sample buffer for 10 min at 95 °C. For MYC-IP, 3 mg of cell lysate was incubated with 20 µL of MYC-trap bead slurry in dilution buffer, 10 mM Tris-HCl (pH 7.5), 150 mM NaCl, 0.5 mM EDTA, and 1x of PIC for 3 h at 4 °C in an end-over-end rotator. After IP incubation, the beads were centrifuged for 2 min at 2,500 × g, 4 °C and were washed with dilution buffer three times. At the end, proteins were eluted from the beads by heating the beads in 2x SDS sample buffer at 95 °C for 10 min.

#### Immunofluorescence and microscopy

For staining free cholesterol with LAMP2 staining, MEF and HeLa cells were seeded in 24-well plates at the density of 8 x 10^3^ and 15 x 10^4^ cells per well, respectively. Cells were cultured for 24 h after seeding and used for staining. For staining, cells were washed with PBS twice and were fixed with 4% PFA for 10 min at 37 °C. After fixation, the cells were washed with PBS three times and were permeabilized with 0.1% Triton for 10 min at RT. Subsequently, the cells were incubated with 125 µg/mL of filipin for 2 h at RT. Next, the cells were washed three times with PBS and were blocked with 10% FCS in PBS for 1 h at RT. After blocking, the cells were incubated with the anti-LAMP2 primary antibody overnight in a humid chamber. Next day, the cells were washed with TBS three times, 5 min each. The cells were incubated in the respective secondary antibody for 1 h at RT in dark. After secondary antibody incubation, the cells were washed three times with TBS for 5 min each, rinsed once with distilled water, mounted on cover slides using ProLongTM Gold antifade mounting medium. The cover slides were dried overnight at RT before imaging.

For other immunostaining, MEF and HeLa cells were seeded in 12-well plates at the density of 1 x 10^4^ and 25 x 10^4^ cells per well, respectively. Cells were cultured for 24 h after seeding and used for staining. For staining, cells were washed with PBS twice and were fixed with 4 % PFA for 10 min at 37 °C. After fixation, the cells were washed with PBS three times and were permeabilized with 0.1% Triton for 10 min at RT. Next, the cells were washed three times with PBS and were blocked with 10 % FCS in PBS for 1 h at RT. After blocking, the cells were incubated with the respective primary antibodies overnight in a humid chamber. Next day, the cells were washed with TBS three times, 5 min each. The cells were incubated in the respective secondary antibodies for 1 h at RT in dark. After secondary antibody incubation, the cells were washed three times with TBS for 5 min each, rinsed once with distilled water, mounted on cover slides using ROTI Mount FluorCare DAPI. The cover slides were dried overnight before imaging.

Images from MEF, HeLa, and U2OS U18666A-treated cells were captured using Leica SP5 AOBS with SMD confocal microscope equipped with an HCX PL APO×63/oil objective and 2x single-photon avalanche diode detector. Images were acquired using the LAS AF software. Images were annotated using Fiji software.^87^ Colocalization and lysosome positioning analyses were performed as per Schmied et al.^88^ Images from ARL8B and BORCS5 siRNA experiments were captured using an Axiovert 200 M microscope equipped with an AxioCam705 camera; images from HA-LAMTOR1-LLL-AAA experiments were captured using a Zeiss Axio Observer.Z1/7 Airyscan microscope. These images were acquired using the ZEN software 3.4.

#### Acetyl-Coenzyme A assays

The Acetyl-Coenzyme A assay (#MAK039) was performed as per the manufacturer instructions.

#### Radioactive protein kinase screening assay

The radioactive kinase testing including 245 serine/threonine kinases was performed by Reaction Biology Europe GmbH, Freiburg, Germany. The assay was conducted with and without biotinylated LAMTOR1 peptide (DEQALLSSILAKTA). The assay plates were incubated at 30 °C for 60 min. Subsequently, the reaction was stopped using 4.7 M NaCl/ 35 mM EDTA. The reaction mixture was transferred to 96-well streptavidin-coated ScintiPlates followed by 30 min incubation at RT. The plates shaken to allow for binding of the biotinylated peptides to the streptavidin. Next, the plates were aspirated and washed using 0.9% NaCl. The incorporation of radioactive 33Pi was determined using a microplate scintillation counter.

#### *In vitro* kinase assay

*In vitro* kinase reactions were performed in kinase buffer consisting of 25 mM HEPES (pH 7.4), 75 mM NaCl, 0.9 mM TCEP and 5% glycerol. Reactions were carried out at 30 °C for 4 min using the hyperactive mTORC1 composite mutant (T1977R + S2215Y on mTOR) (samples provided by Madhanagopal Anandapadamanaban, Roger Williams Lab, MRC Laboratory of Molecular Biology, Cambridge, UK). Reactions were set up by pre-incubating mTORC1 with RagA/C and Ragulator complex for 10 min on ice. Samples were then equilibrated at 30 °C for 15 s, and kinase assays were initiated by the addition of ATP (250 µM) and MgCl₂ (10 mM) to final concentrations. Reactions were stopped by adding EDTA to a final concentration of 50 mM.

#### Cell perturbations

##### Preparation of LPDS

The density of FCS was adjusted to 1.21 g/mL using solid KBr and the density-adjusted FCS was centrifuged for 24 h at 20,000 × g. After centrifugation, the lipoprotein-free serum was aspirated carefully using a syringe-needle. Next, the lipoprotein-free serum was dialyzed using the dialysis buffer containing 150 mM NaCl, 10 mM Tris, and 2 mM EDTA (pH 8.6) at 4 °C.

##### LPDS treatment

1 x 10^6^ cells were seeded 24 h prior to the treatment. The medium was removed and the cells were rinsed with PBS twice. Next, the cells were incubated with starvation medium for 17 h at 37 °C.

##### MCD treatment

For HeLa cells, 1 x 10^6^ cells were seeded per 10 cm dish 24 h prior treatment. On the day of treatment, cells were washed twice with serum-free media, supplemented with serum-free medium containing 0.5% LPDS and 0.5% of MCD, and were incubated for 2 h at 37 °C.

##### Amino acid starvation

Cells were seeded to obtain 80% confluence 24 h prior to the treatment. On the day of treatment, cells were washed three times with amino acid-free media (7.84 g DMEM Ham’s F-12 w/HEPES, NaHCO_3_, without amino acids in 400 mL of water, pH 7.2.) containing 10% dialyzed FCS, and were incubated for 5 h at 37 °C.

##### U18666A treatment

Cells were supplemented with 3 µg/mL of U18666A and were incubated for 24 h at 37 °C.

##### OSW-1 treatment

Cells were supplemented with 200 nM of OSW-1 and were incubated for 24 h at 37 °C.

#### Thin layer chromatography

Silica gel plates were activated for 5 min at 180 °C. Samples were dissolved in CHCl_3_ and spotted onto silica gel plates using a micropipette. The samples were developed in a mobile phase in a vertical closed chamber. After development, the plates were air dried and were sprayed using CuSO_4_/H3PO_4_. Subsequently, the plates were incubated at 180 °C for 7 min.

#### Quantification and statistical analysis of western blots

Data are plotted as mean ± SD or mean ± SEM as mentioned in the figure legends. The samples sizes are indicated as datapoints or mentioned in the figure legends. The number of replicates is also indicated in the figure legends. Statistical analysis was performed using Student’s t-test, or one-way ANOVA. All the statistical analyses were performed using Graphpad Prism 6.01. Statistical significance is denoted as * p < 0.05, ** p < 0.01, and *** p < 0.001.

## Abbreviations

ACC: Acetyl-CoA carboxylase
AKT: Protein kinase B
AMPK: 5’-AMP activated protein kinase
APS: Ammonium persulfate
ARL8B: ADP-ribosylation factor-like protein 8B
ATP: Adenosine triphosphate
BORC: BLOC-one-related complex
BORCS5: BORC subunit 5
BORCS6: BORC subunit 6
BSA: Bovine serum albumin
co-IP: co-immunoprecipitation
CREB: cAMP response element binding protein
crRNA: CRISPR RNA
DDA: Data-dependent acquisition
DMEM: Dulbecco’s modified Eagle’s media
DMSO: Dimethylsulfoxide
DTT: Dithiothreitol
ER: Endoplasmic reticulum
FA: Formic acid
FCS: Fetal calf serum
FKBP12: Peptidyl-prolyl cis-trans isomerase FKBP1A
GO: Gene ontology
GSEA: Gene set enrichment analyses
GTP: Guanosine triphosphate
HEPES: 2-[4-(2-Hydroxyethyl)-1-piperazinyl]-ethanesulfonic acid
JIP4: C-Jun-amino-terminal kinase-interacting protein 4
KCl: Potassium chloride
KH_2_PO_4_: Monopotassium phosphate
LAMTOR: Late endosomal and lysosomal adaptor and MAPK
LDL: Low-density lipoprotein
LEF: Lysosome-enriched fraction
LIPA: Lysosomal acid lipase
LLL: Lysosomal sorting motif
LLOMe: L-leucyl-L-leucine methyl ester hydrobromide
LPDS: Lipid-depleted serum
MAP4K2: Mitogen-activated protein kinase kinase kinase kinase 2
MAPK: Mitogen-activated protein kinase
MCD: Methyl-β-cyclodextrin
MEF: Mouse embryonic fibroblasts
MeOH: Methanol
MS: Mass spectrometry
mTOR: Mechanistic target of rapamycin
NH_4_HCO_3_: Ammonium bicarbonate
NPC1/2: Niemann Pick Type C 1/2
ORP1L: OSBP-related protein 1L
OSBPs: Oxysterol-binding proteins
P70 S6K: 70-kilodalton ribosomal protein S6 kinase
PAGE: Polyacrylamide gel electrophoresis
PBS: Dulbecco’s phosphate-buffered saline
PCR: Polymerase chain reaction
PFA: Paraformaldehyde
PhosSTOP: Phosphatase inhibitor cocktail
PI(4)P: Phosphatidylinositol 4-phosphate
PIC: Protease inhibitor cocktail
PKA: Protein kinase A
PLL: Poly-L-Lysine
PNS: Post-nuclear supernatant
RT: Room temperature
S56: Serine 56
S6 RP: S6 ribosomal protein
SCARB2/LIMP2: Lysosomal integral membrane protein 2
SDS: Sodium dodecyl sulfate
SILAC: Stable isotope labeling by amino acids in cell culture
SKIP: Pleckstrin homology domain-containing family M member 2
SPIONs: Superparamagnetic iron oxide nanoparticles
SREBP1/2: Sterol regulatory element-binding proteins 1 and 2
TBST: Tris-buffered saline containing 0.1% Tween 20
TEAB: Triethylammonium bicarbonate
TEMED: Tetramethylethylenediamine
TFA: Trifluoroacetic acid
TFEB: Transcription factor EB
TLC: Thin layer chromatography
TMEM55B: Lysosomal transmembrane protein 55B
U2OS: Human Osteosarcoma
VAPA/B: Vesicle-associated membrane protein-associated proteins A and B
v-ATPase: vacuolar-type ATPase
WCL: Whole cell lysate

---

**Figure S1.**
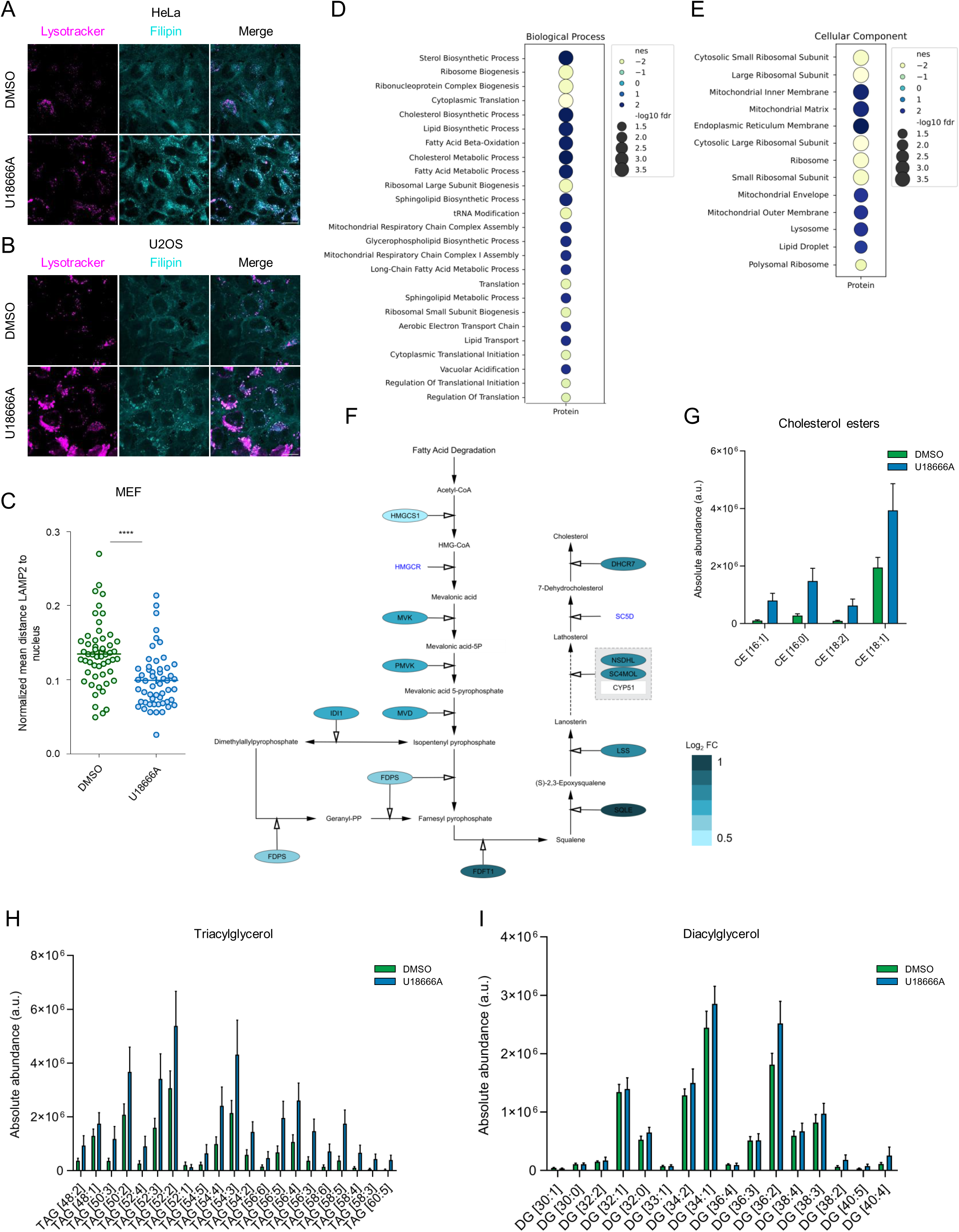
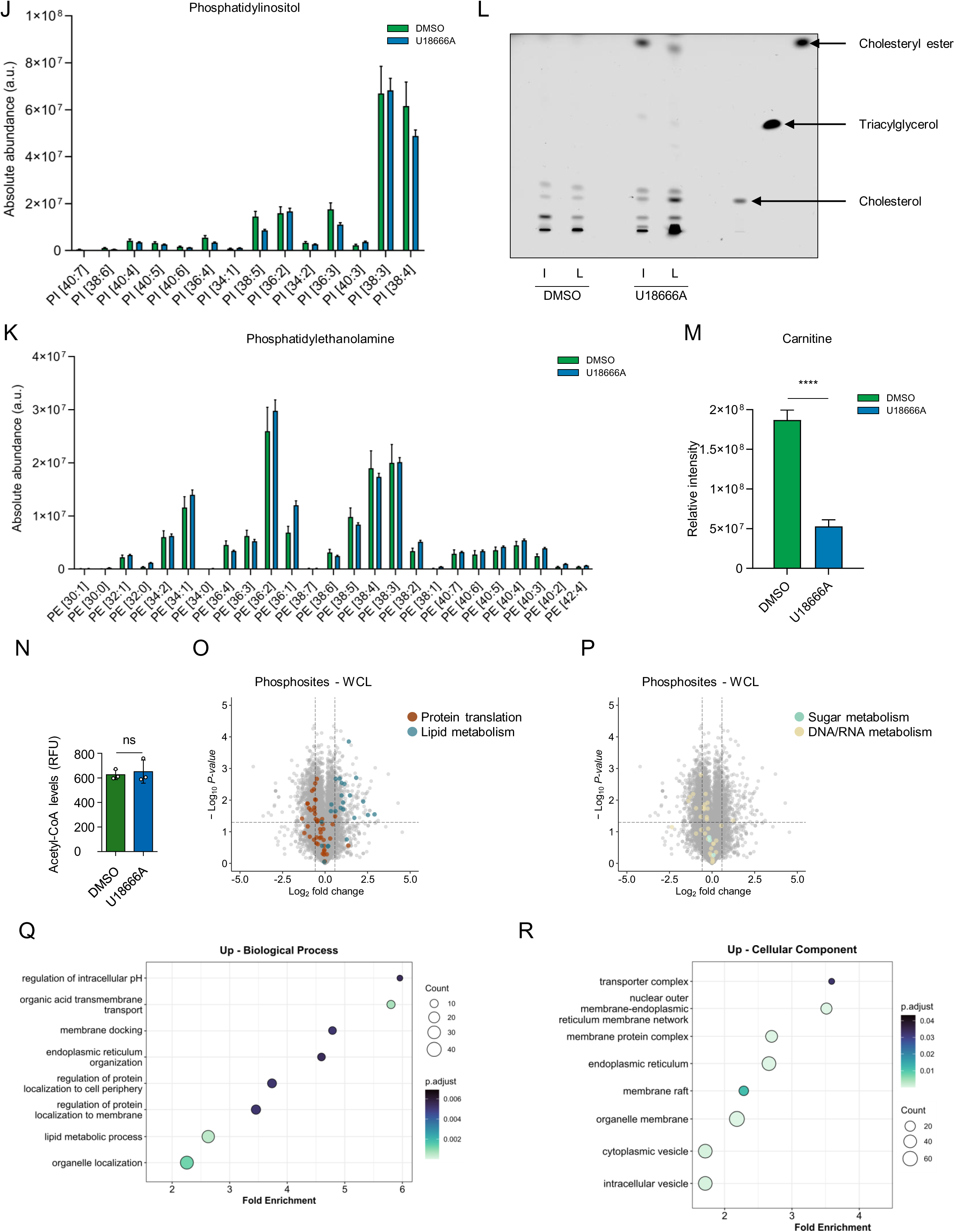
Multi-OMICs analysis of impaired lysosomal cholesterol egress identifies regulatory patterns on the protein, phosphorylation site, and lipid level (related to Figure 1). (A) HeLa and (B) U2OS cells were treated with U18666A (3 µg/mL) or DMSO (0.03%) as a vehicle control for 24 h followed by staining using lysotracker and Filipin and subsequently imaged by confocal microscopy. Scale bar, 10 µm. (C) In Filipin-stained MEF cells, nuclei were manually drawn using Fiji to facilitate the measurement of lysosomal distance to the nucleus. Plotted are the normalized mean distance of LAMP2/Filipin stained punctae on the y-axis for individual cells (n = 46). Data represented are mean ± SEM. Statistical significance was determined using unpaired two-tailed t-test. **** p < 0.0001. (D, E) Dot plot showing selected gene set enrichment analysis (GSEA) results of U18666A/DMSO-treated MEF cells for the category (D) biological process and (E) cellular components. Nes, normalized enrichment score. For all the results, see Supplemental Table 2. (F) Proteomic data were mapped onto the cholesterol biosynthetic-reactome pathway using Cytoscape. Nodes represent proteins, color-coded by log_2_ fold change (dark blue, upregulated; light blue, downregulated). (G, H, I, J, K) Bar graphs showing levels of lipid species of (G) cholesteryl esters (CE), (H) triacylglycerols (TAG), (I) diacylglycerols (DG), (J) phosphatidylinositols (PI), (K) phosphatidylethanolamines (PE) measured by LC-MS/MS in U18666A-vs DMSO-treated MEF cells. Data are presented as mean ± SD (n = 5 biological replicates). (L) Thin layer chromatography of WCL (input, I) and LEF (lysosome, L) from U18666A- and DMSO- treated MEF cells. (M) Bar graph showing relative intensities of carnitines in U18666A-vs DMSO-treated MEF cells. Data are presented as mean ± SD (n = 5 biological replicates). Statistical significance was determined using unpaired two-tailed t-test. **** p < 0.0001. (N) Acetyl-CoA levels were quantified using the Acetyl-CoA assay kit in DMSO- and U18666A-treated MEF cells. Levels are expressed in relative fluorescence unit (RFU). Data presented as mean ± SD (n = 3 biological replicates). Statistical significance was determined using unpaired two-tailed t-test, ns: non-significant. (O) Volcano plot depicting changes in protein phosphorylation levels in WCL, as determined by phosphoproteomic analysis of U18666A-treated versus DMSO-treated MEF cells. Differentially regulated protein phosphorylation sites involved in protein translation and cholesterol metabolism, manually curated from the literature are highlighted. (P) Volcano plot depicting changes in protein phosphorylation levels in WCL, as determined by phosphoproteomic analysis of U18666A-treated versus DMSO-treated MEF cells. Differentially regulated protein phosphorylation sites involved in sugar and nucleic acid metabolism, manually curated from the literature are highlighted. (Q, R) Dot plot showing selected significantly enriched gene ontology (GO) terms for the category (R) abiological processs and (S) cellular component in U18666A/DMSO-treated MEF cells. For all details see Supplemental Table 2.

**Figure S2.**
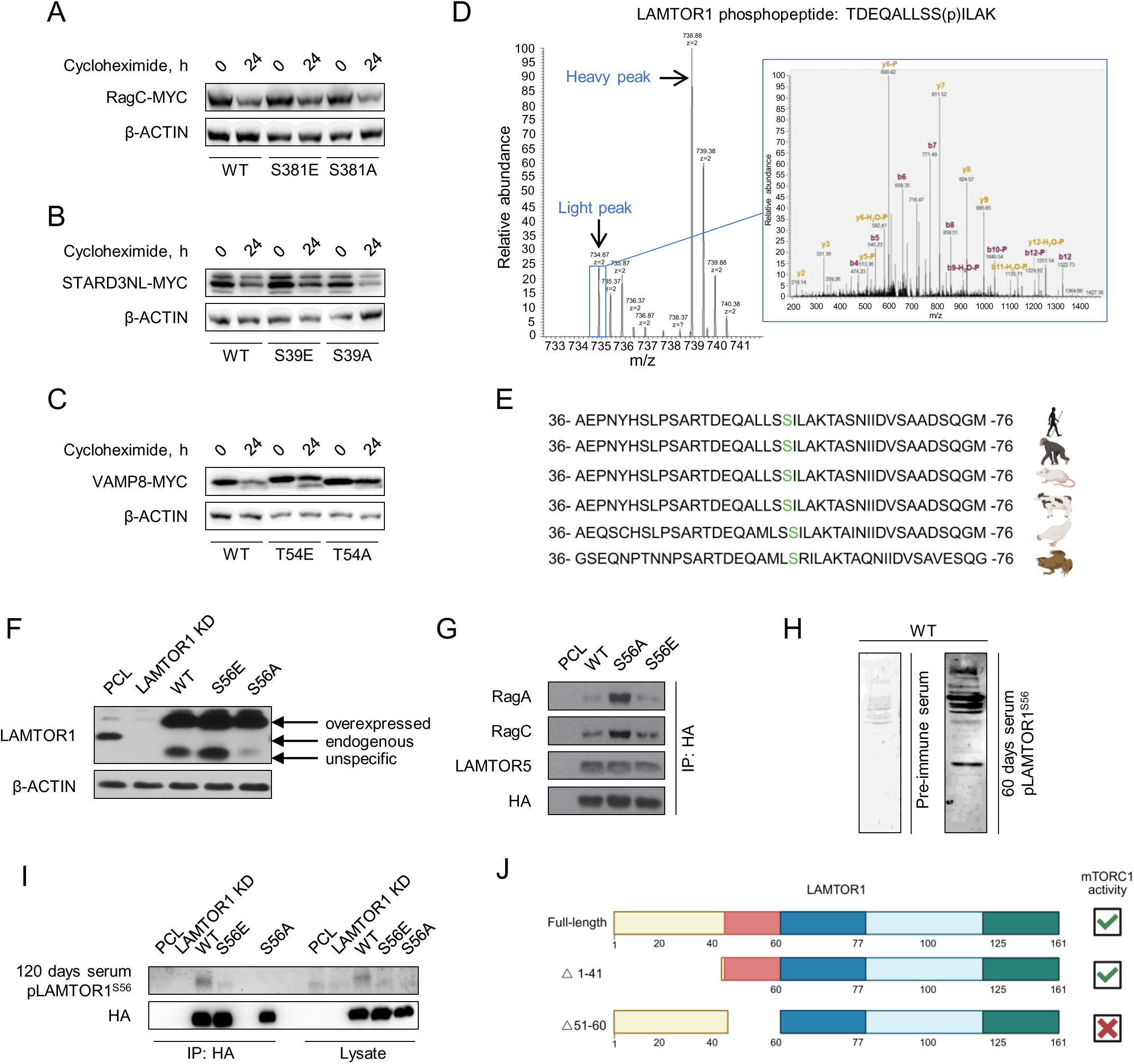
Generation of stable cells and antibodies for investigation of LAMTOR1 pS56 (related to Figure 2). (A, B, C) HEK293 cells were transfected with MYC-tagged WT, E, or A versions of RagC, STARD3NL, or VAMP8 followed by incubation with cycloheximide for the specified periods. Cell lysates were probed with MYC antibody. (D) Manually annotated MS/MS spectrum of LAMTOR1 pS56, with fragment ions (b- and y-ions) labeled to show the fragmentation pattern. (E) Sequence alignment of LAMTOR1 pS56 peptide across different species, with conserved residues highlighted in color. (F) Generation of HA-tagged LAMTOR1 WT, or S56A cell lines in a LAMTOR1 knockdown background. Cell lysates from PCL, LAMTOR1 knockdown (KD), WT, S56E, or S56A were probed with LAMTOR1 antibody and loading control. (G) Rag GTPase binding to Ragulator is regulated by LAMTOR1 S56 phosphorylation status. Lysates from HA-tagged LAMTOR1 WT, S56E, or S56A cells were subjected to HA-IP, and IP eluates were probed with antibodies indicated. (H) Verification of affinity purified LAMTOR1 phosphosite specific antibody. Cell lysates from HA-tagged LAMTOR1 WT were probed with pre-immune serum and anti-serum collected 60 days post-immunization using LAMTOR1 pS56 peptide. (I) Cell lysates from PCL, LAMTOR1 KD, HA-tagged LAMTOR1 WT, S56E, or S56A were subjected to HA-IP, and IP eluates were analyzed for the phosphorylation status of LAMTOR1 S56 and total protein abundance. (J) U18666A kinetics in different versions of LAMTOR1. HA-tagged LAMTOR1 WT, S56E, or S56A were treated with U18666A (3 µg/mL) for the indicated time period. Cell lysates were analyzed for the phosphorylation status of P70 S6K T389 and S6 RP S240/244 using immunoblotting. (K) Dependence of mTORC1 activation on LAMTOR1 Regions. Deletion of the amino acid sequence harboring S56 results in mTORC1 deactivation.^51^

**Figure S3.**
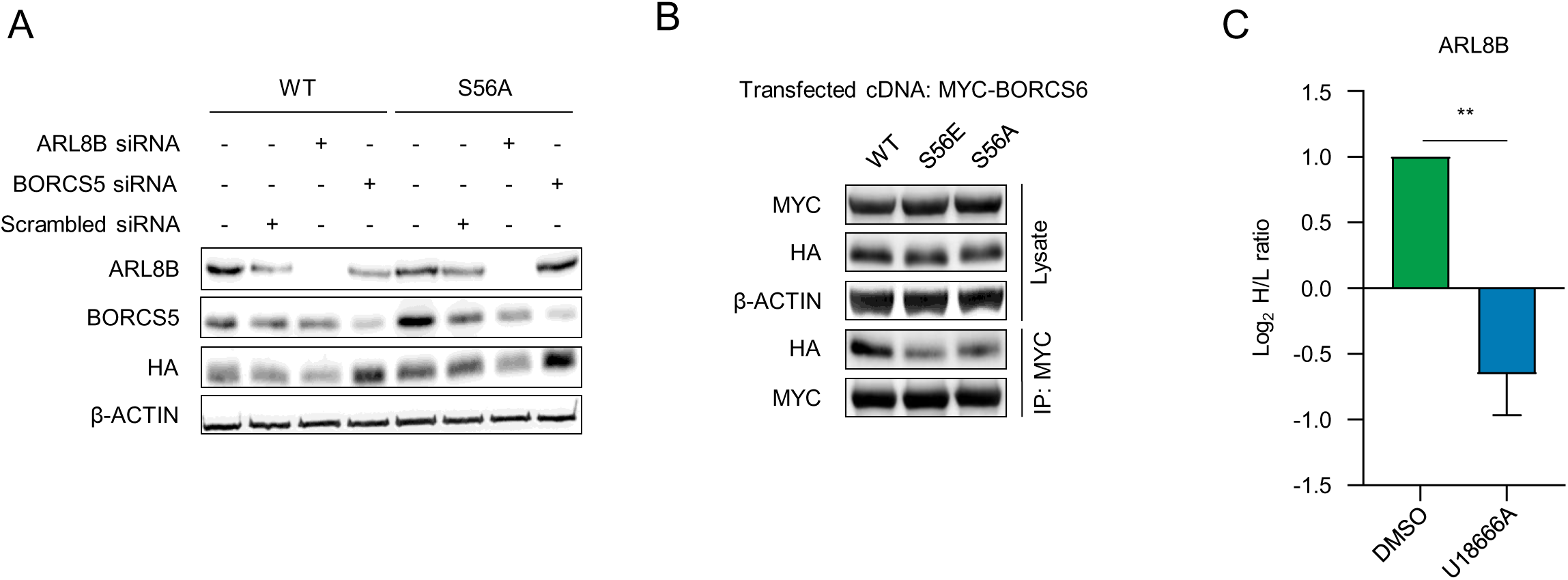
Investigation of LAMTOR1 pS56’s effect on BORC (related to Figure 3). (A) Validation of siRNA-mediated knockdown of ARL8B and BORCS5. HA-tagged LAMTOR1 WT, or S56 cells were transfected with scrambled, ARL8B or BORCS5 siRNAs. Cell lysates were probed with antibodies targeting ARL8B, BORCS5, HA, or actin. (B) HA-tagged LAMTOR1 WT, S56E and S56A cells were transfected with MYC-BORCS6 cDNA followed by MYC-IP. Left: Cell lysates and IP eluates were probed with antibodies, as indicated. (C) Differential regulation of ARL8B in LEF in response to lysosomal cholesterol accumulation. Log2 fold change (U18666A versus DMSO) shown as mean ± SD (n = 3 biological replicates). Statistical significance was determined using unpaired two-tailed t-test. ** p < 0.01.

**Figure S4.**
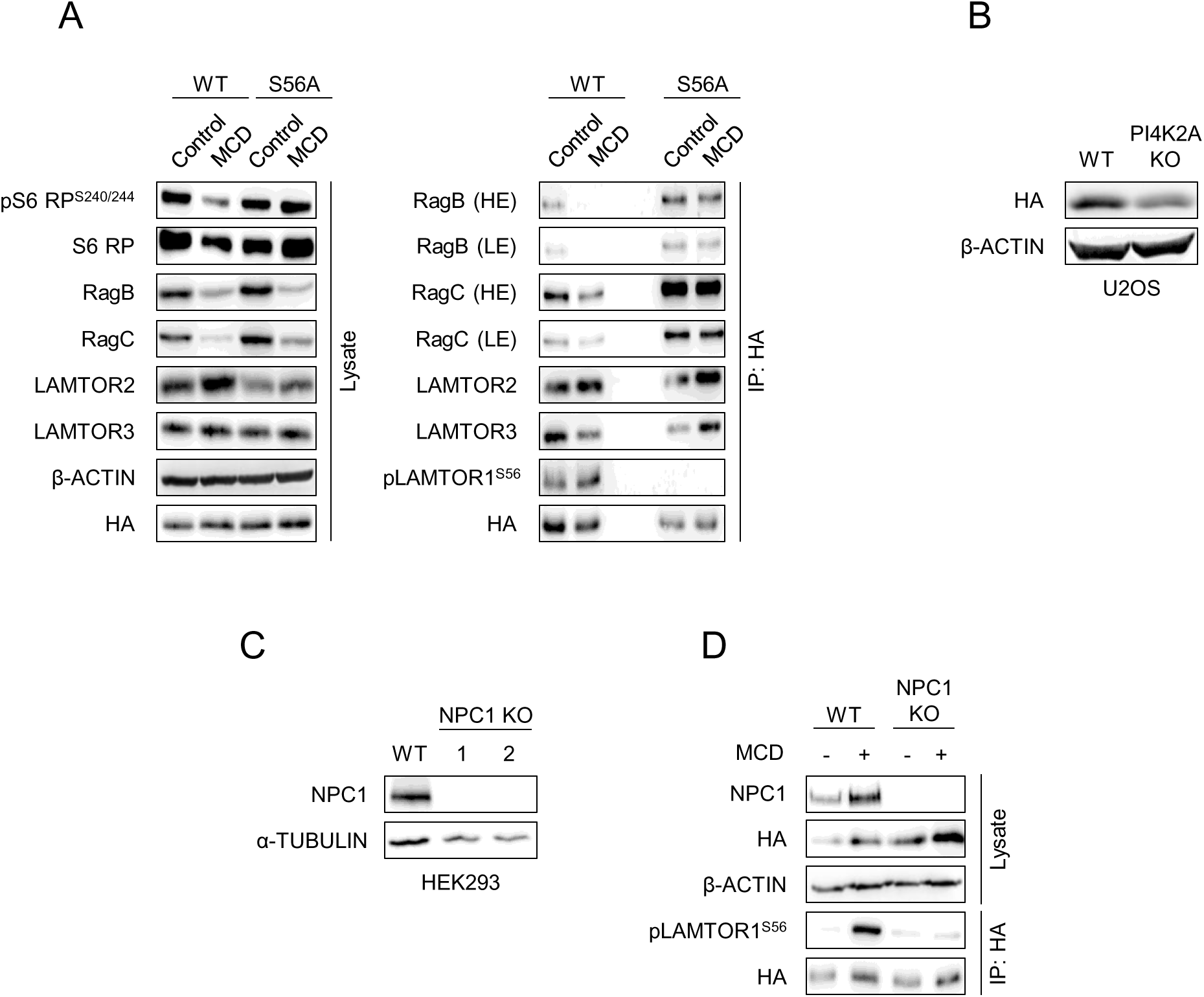
Characterization of mechanisms regulating LAMTOR1 S56 phosphorylation (related to Figure 4). (A) LAMTOR1 phosphorylation induced by depletion of membrane cholesterol levels displaces Rag GTPases. HA-tagged LAMTOR1 WT or S56A cells were treated with 0.5% MCS for 2 h in combination with 3% LPDS. Cell lysates were subjected to HA-IP. Both IP eluates and cell lysates were probed with antibodies as indicated. (B) Validation of PI4K2A KO cells expressing HA-tagged LAMTOR1 WT. (C) Validation of CRISPR-Cas9 generated HEK293 NPC1 KO cells. (D) HEK293 NPC1 KO cells expressing HA-tagged LAMTOR1 WT were treated with 0.5% MCD in combination with 3% LPDS for 2 h followed by HA-IP. Immunoprecipitates were used to monitor phosphorylation status of LAMTOR1 S56 phosphorylation by immunoblotting.

**Figure S5.**
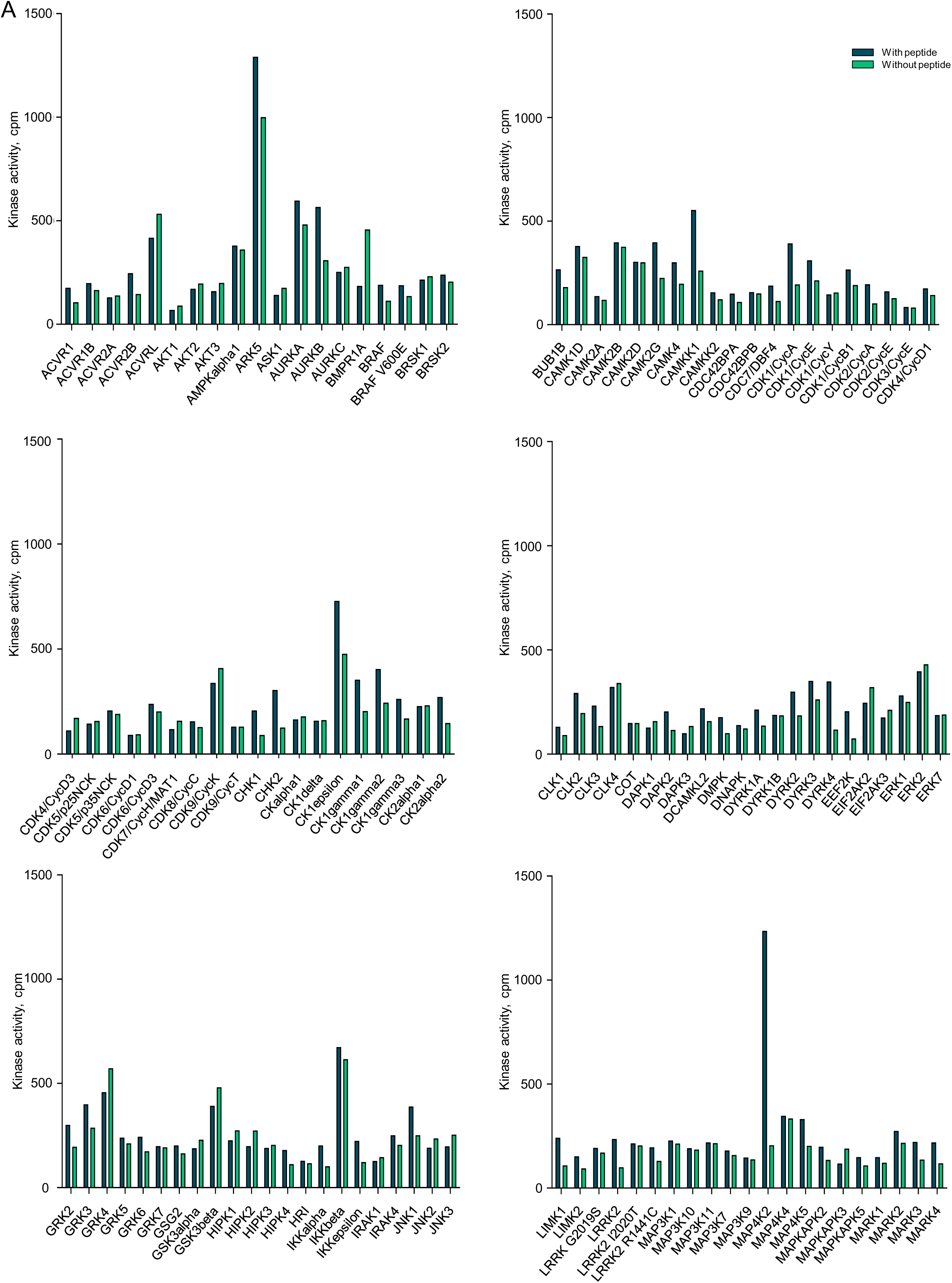
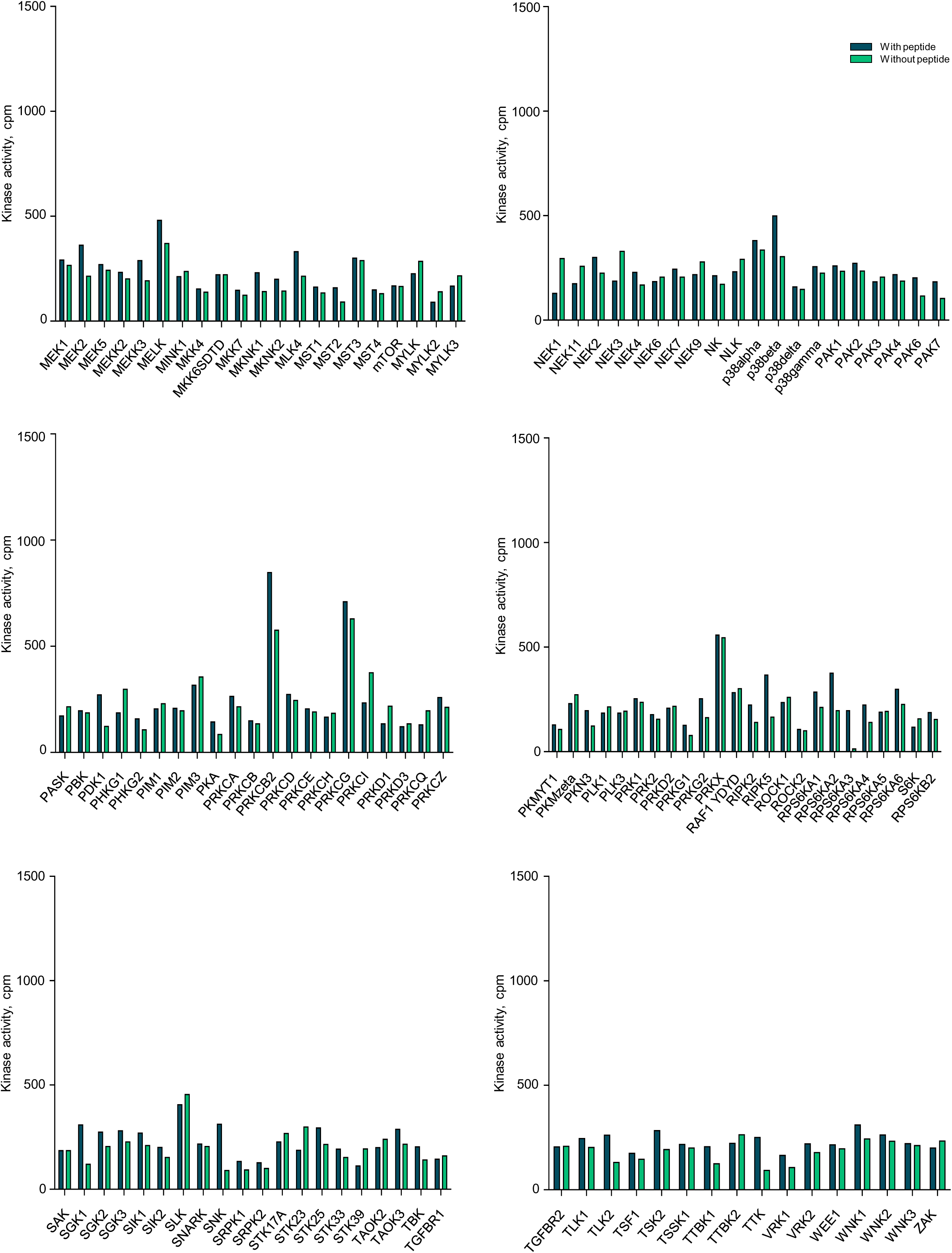
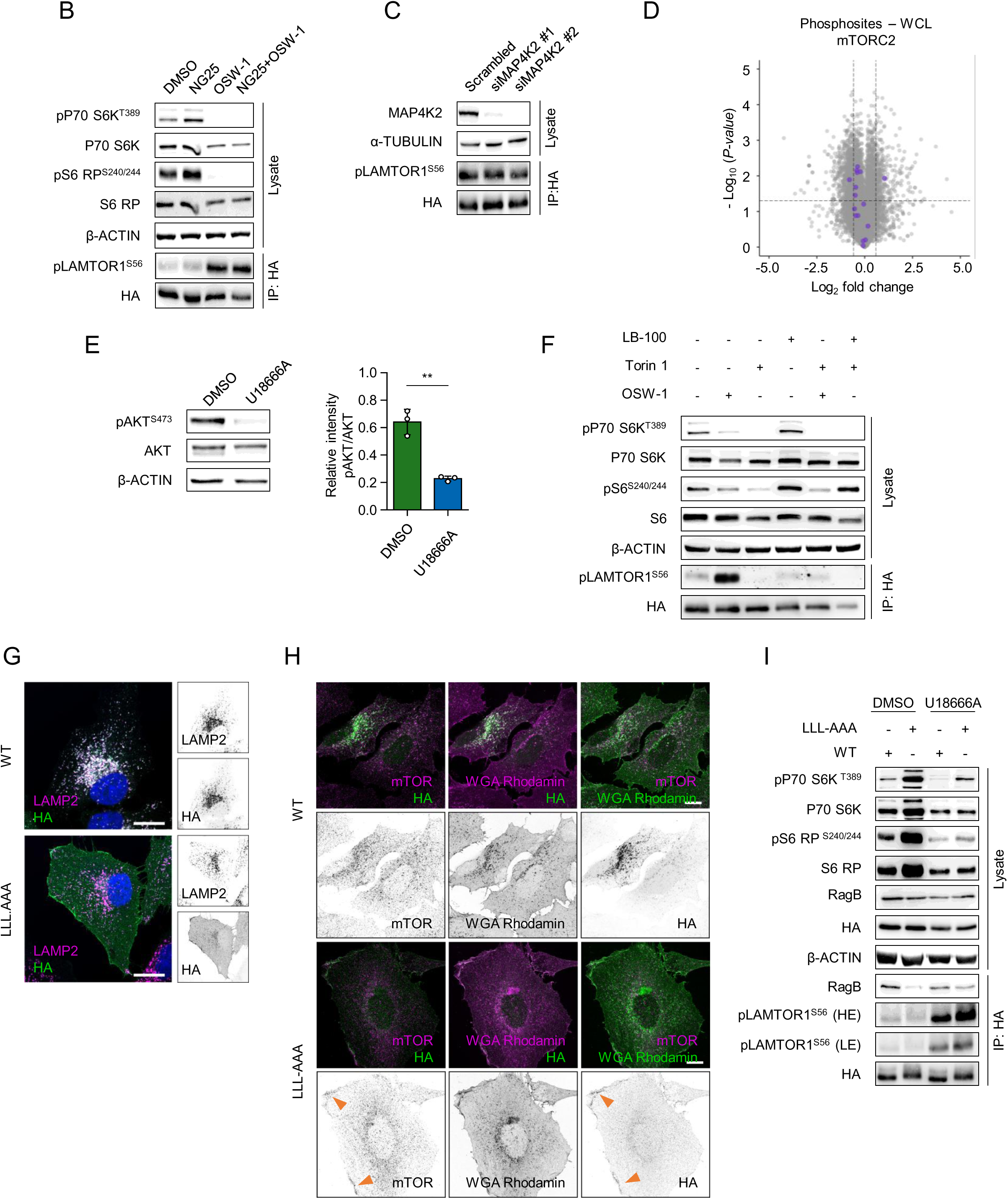
Investigation of the kinase phosphorylating LAMTOR1 S56 (related to Figure 5). (A) 245 serine/threonine kinases were individually incubated with or without biotinylated LAMTOR1 peptide (DEQALLSSILAKTA) and kinase activity was measured using a radioactive ATP assay. (B) MAP4K2 inhibition does not affect LAMTOR1 phosphorylation. HA-tagged LAMTOR1 WT cells were treated with 1 µM NG25, 200 nM OSW-1 or a combination of both for 2 h followed by HA-IP. Cell lysates and IP eluates were subjected to immunoblotting. Phosphorylation status of LAMTOR1 S56 was monitored using the LAMTOR1 phosphosite antibody. (C) MAP4K2 silencing does not alter LAMTOR1 phosphorylation. HA-tagged LAMTOR1 WT cells were transfected with scrambled, MAP4K2 #1 or MAP4K2 #2 siRNAs followed by HA-IP. Cell lysates and IP eluates were subjected to immunoblotting. Phosphorylation status of LAMTOR1 S56 was monitored using LAMTOR1 phosphosite antibody. (D) U18666A downregulated phosphorylation levels of mTORC2 substrates. Volcano plot depicting changes in phosphorylation levels of mTORC1 substrates in WCL, as determined by phosphoproteomic analysis of U18666A-treated versus DMSO-treated MEF cells. (E) mTORC2 may be unrelated to LAMTOR1 phosphorylation. Left: MEF cells were treated with U18666A (3 µg/mL) for 24 h and cell lysates were probed with phosphosite specific antibodies against a well-known mTORC2 substrate, AKT. Right: Densitometric quantification of immunoblots and data shown are mean ± SD (n=3 biological replicates, indicated by data points) of relative band intensities. Statistical significance was determined using unpaired two-tailed t-test. Asterisks denote statistical significance as follows: ** p < 0.01. (F) HA-tagged LAMTOR1 WT cells were treated with 250 nM Torin 1 for 50 min, 200 nM OSW-1 for 2 h, 2 µM LB-100 (PP2A inhibitor) for 3 h or combination of these compounds as indicated. IP eluates and cell lysates were probed with antibodies as indicated. (G) Validation of plasma-membrane localized LAMTOR1. HA-tagged LAMTOR1 WT or LLL-AAA (plasma membrane-localization mutant) expressing cells were stained using LAMP2 and HA antibody, and imaged by confocal microscopy. Magnified inset shown at the side. Scale bar, 10 µm. (H) HA-tagged LAMTOR1 WT or LLL-AAA expressing cells were stained using WGA Rhodamin, LAMP2 antibody and HA antibody in different combinations as indicated. The cells were subsequently imaged by confocal microscopy. Scale bar, 10 µm. (I) HA-tagged LAMTOR1 WT or LLL-AAA cells were treated with U18666A (3 µg/mL) for 24 h followed by HA-IP. IP eluates and cell lysates were subjected to immunoblotting using the indicated antibodies. Phosphorylation levels of all proteins detected by phospho-specific antibodies in immunoblots were quantified and normalized to their corresponding total protein levels.

